# Pptc7 maintains mitochondrial protein content by suppressing receptor-mediated mitophagy

**DOI:** 10.1101/2023.02.28.530351

**Authors:** Natalie M. Niemi, Lia R. Serrano, Laura K. Muehlbauer, Catie Balnis, Keri-Lyn Kozul, Edrees H. Rashan, Evgenia Shishkova, Kathryn L. Schueler, Mark P. Keller, Alan D. Attie, Julia Pagan, Joshua J. Coon, David J. Pagliarini

## Abstract

Pptc7 is a resident mitochondrial phosphatase essential for maintaining proper mitochondrial content and function. Newborn mice lacking *Pptc7* exhibit aberrant mitochondrial protein phosphorylation, suffer from a range of metabolic defects, and fail to survive beyond one day after birth. Using an inducible knockout model, we reveal that loss of *Pptc7* in adult mice causes marked reduction in mitochondrial mass concomitant with elevation of the mitophagy receptors Bnip3 and Nix. Consistently, *Pptc7*^-/-^ mouse embryonic fibroblasts (MEFs) exhibit a major increase in mitophagy that is reversed upon deletion of these receptors. Our phosphoproteomics analyses reveal a common set of elevated phosphosites between perinatal tissues, adult liver, and MEFs— including multiple sites on Bnip3 and Nix. These data suggest that *Pptc7* deletion causes mitochondrial dysfunction via dysregulation of several metabolic pathways and that Pptc7 may directly regulate mitophagy receptor function or stability. Overall, our work reveals a significant role for Pptc7 in the mitophagic response and furthers the growing notion that management of mitochondrial protein phosphorylation is essential for ensuring proper organelle content and function.

## Introduction

Mitochondria engage in numerous cellular processes in eukaryotic cells, including ATP synthesis via oxidative phosphorylation, nutrient selection, energy expenditure, cofactor biosynthesis, and the commitment to cell death. Over time, mitochondria can become damaged, requiring quality control mechanisms to maintain a healthy mitochondrial population (Song et al., 2021). One such mechanism is mitophagy, which engages the autophagic machinery to clear damaged or superfluous organelles (Pickles et al., 2018). A growing body of work suggests that protein phosphorylation plays a key role in regulating mitophagy (Wang et al., 2020), and in various other mitochondrial processes such as protein import (Kravic et al., 2018; Walter et al., 2021), core metabolism (Grimsrud et al., 2012; Guo et al., 2017), heme biosynthesis (Chung et al., 2017), and programmed cell death (Niemi & MacKeigan, 2013). Indeed, multiple candidate protein phosphatases reside at or within mitochondria (Calvo et al., 2016; Niemi & Pagliarini, 2021; Pagliarini et al., 2008; Rath et al., 2021) where they would be positioned to manage phosphorylation on mitochondrial proteins. Despite their prevalence, many of these phosphatases remain poorly studied.

Our recent work demonstrates that loss of one phosphatase, Pptc7, leads to the hyperphosphorylation of select mitochondrial proteins and significant loss of overall mitochondrial protein content (Niemi et al., 2019). *Pptc7* knockout (KO) animals present with hypoketotic hypoglycemia and fail to survive a single day after birth. *Pptc7* knockout tissues show widespread decreases in mitochondrial protein levels with little to no effects on matched mRNA levels, suggesting altered post-transcriptional control of mitochondrial content.

To further explore this phenomenon, we generated two independently derived *Pptc7* knockout models: a conditional mouse that allows inducible *Pptc7* knockout in adult mice, and isolated fibroblasts from our global *Pptc7* (i.e., perinatal lethal) knockout mouse model. We find that knockout of *Pptc7* decreases mitochondrial protein content in all tested systems. Interestingly, the mitophagy receptors Bnip3 and Nix were among the limited number of proteins that increased in expression across both mouse models and isolated fibroblasts. Our recent MITOMICS study revealed that *PPTC7* loss also drives elevation of these receptors in human cells, and this response was unique across over 200 knockout cell lines (Rensvold et al., 2022). CRISPR-mediated knockout of Bnip3 and Nix (gene name *Bnip3l*) rescued the expression of most mitochondrial proteins in *Pptc7* KO cells, demonstrating that these receptors are necessary for the observed mitophagy. However, mitochondrial dysfunction persisted in *Pptc7* knockout cells after *Bnip3* and *Bnip3l* knockout, indicating that this dysfunction is connected at least partially to the underlying aberrant protein phosphorylation. Our phosphoproteomic analyses suggest a varied set of substrates for the phosphatase Pptc7, including the import protein Timm50, multiple core metabolic proteins, and the Bnip3 and Nix mitophagy receptors themselves. Collectively, these data demonstrate that Pptc7 is critical for managing mitochondrial protein phosphorylation as well as regulating Bnip3- and Nix-associated mitophagy.

## Results

### Pptc7 maintains mitochondrial protein levels in adult mice

We demonstrated previously that knockout (KO) of the mitochondrial protein phosphatase *Pptc7* causes decreased mitochondrial content, metabolic dysfunction, and fully penetrant lethality within one day of birth (Niemi et al., 2019). To overcome the limitations associated with these severe phenotypes, we generated a floxed *Pptc7* mouse with the potential for conditional knockout (Figure 1A). We bred floxed *Pptc7* mice to the well-established UBC-Cre-ER^T2^ mouse (Ruzankina et al., 2007) to allow widespread, inducible recombination in response to tamoxifen (Figure 1B). We validated tamoxifen-induced genomic excision of *Pptc7* exon 3 across seven tissues (Supplementary Figure 1A), which we confirmed at the protein level in liver (Supplementary Figure 1B). Furthermore, age-matched female Cre-positive floxed animals did not show significant levels of recombination in the absence of tamoxifen (Supplemental Figure 1A), indicating that ‘leaky’ Cre-induced knockout of *Pptc7* is not a significant concern through the timeframe of our study.

**Figure 1:**
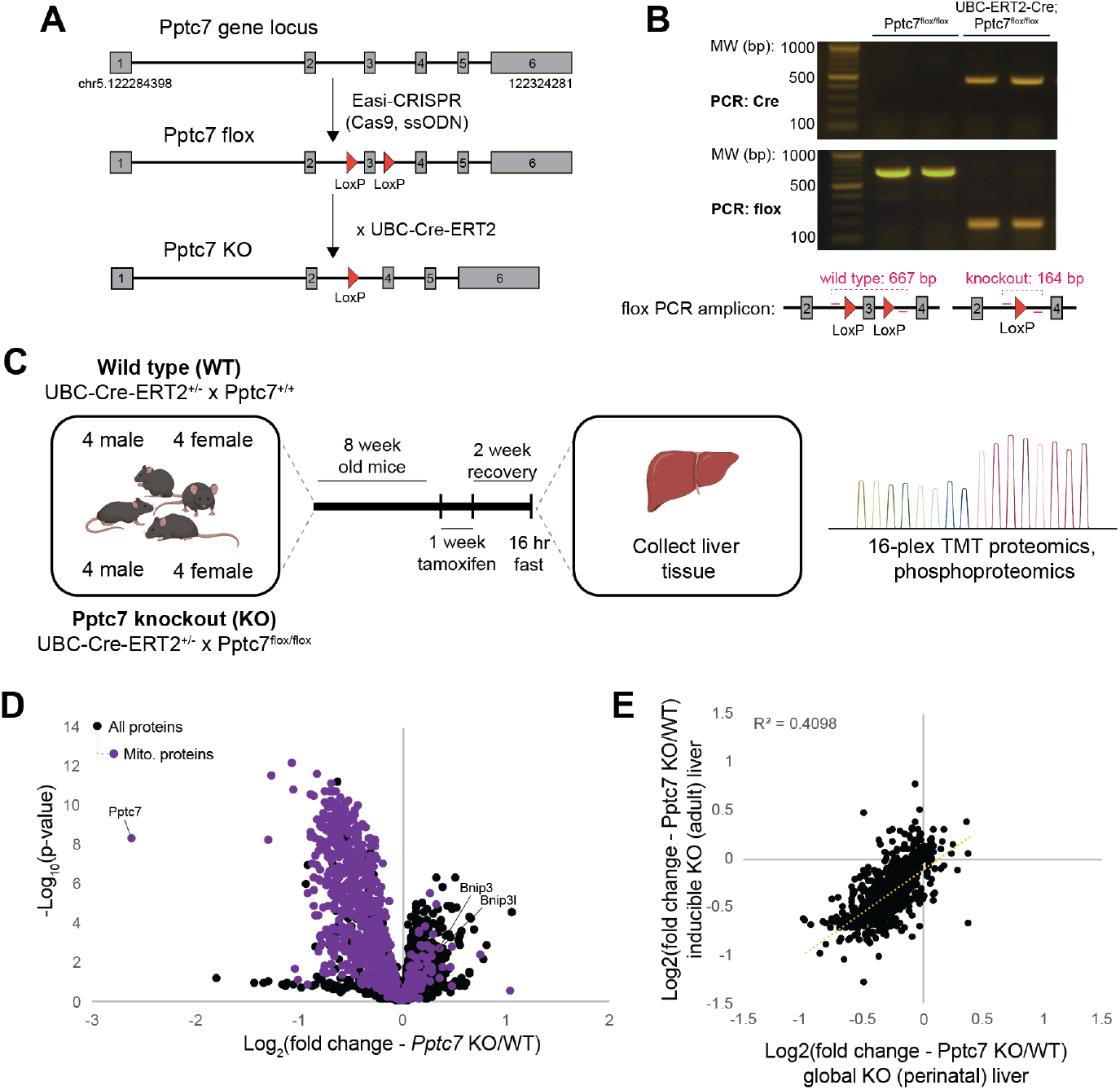
Acute knockout of Pptc7 compromises hepatic mitochondrial content in adult mice. **A**. Schematic for the generation of a conditional *Pptc7* mouse to allow global, inducible knockout in adult mice. **B**. Genotyping verification of Cre-mediated excision of *Pptc7* exon 3 after tamoxifen treatment. **C**. Schematic for the 16plex proteomic analysis of liver tissue from control (UBC-ER^T2^-Cre;Pptc7^+/+^) or experimental (UBC-ER^T2^-Cre;Pptc7^flox/flox^) animals. **D**. Proteomic analysis of non-mitochondrial (black dots) and mitochondrial (purple dots) proteins across 16 liver samples. **E**. Linear regression of the fold changes in mitochondrial proteins identified in both adult inducible KO liver (y-axis) and perinatal KO liver (x-axis).

Upon confirming tamoxifen-inducible KO of *Pptc7* in adult mice, we isolated liver tissue from control (i.e., UBC-Cre^+/-^;Pptc7^+/+^) and knockout (i.e., UBC-Cre^+/-^;Pptc7^flox/flox^) mice of both sexes two weeks post-tamoxifen administration and analyzed these tissues via 16-plex tandem mass tag (TMT) quantitative proteomics (Figure 1C). The results demonstrate that acute loss of *Pptc7* leads to decreased mitochondrial protein content in adult mouse liver comparable to what we observed in the perinatal global *Pptc7* KO model (Figures 1D-E, (Niemi et al., 2019)). Pptc7 was the most significantly downregulated mitochondrial protein in our proteomics analysis (Figure 1D), providing confirmation of *Pptc7* knockout in UBC-Cre^+/-^;Pptc7^flox/flox^ liver tissue. We additionally confirmed decreased mitochondrial content through measurements of citrate synthase (Cs) activity (Supplementary Figure 1C), Cs protein expression (Supplementary Figure 1D), and mitochondrial DNA (mtDNA) levels (Supplementary Figure 1E). Despite the trend of decreased mitochondrial markers, a few proteins, including Bnip3 and its paralog, Nix, were significantly elevated in *Pptc7* KO liver (Figure 1D). An analysis of the proteomic changes between our two KO models revealed significant correlation (Figure 1E), suggesting Pptc7 plays a similar role in liver tissue across developmental states. Collectively, these data demonstrate that acute loss of *Pptc7* in adult mouse liver globally decreases mitochondrial content, suggesting that Pptc7 expression is required across developmental contexts to maintain mitochondrial protein homeostasis.

### Loss of *Pptc7* broadly decreases mitochondrial metabolism in isolated cells

To complement our studies in mice, we established a cell-based system to interrogate mechanisms driving the metabolic and mitochondrial abnormalities in *Pptc7* KO models. We bred *Pptc7*^+/-^ mice from our global KO model to generate mouse embryonic fibroblasts (MEFs) from three wild-type and three knockout E14.5 embryos, which were validated at the gene (Figure 2A) and protein (Figure 2B) levels. A proteomic analysis of these cells revealed significant mitochondrial protein loss (Figure 2C) consistent with our observations from the global *Pptc7* KO heart and liver tissues and inducible *Pptc7* KO liver tissue (although the relative reduction of mitochondrial proteins in vivo is larger than that seen in the MEFs) (Figure 2D). These data demonstrate that loss of *Pptc7* decreases mitochondrial protein content cell autonomously and in a third distinct cell or tissue type.

**Figure 2:**
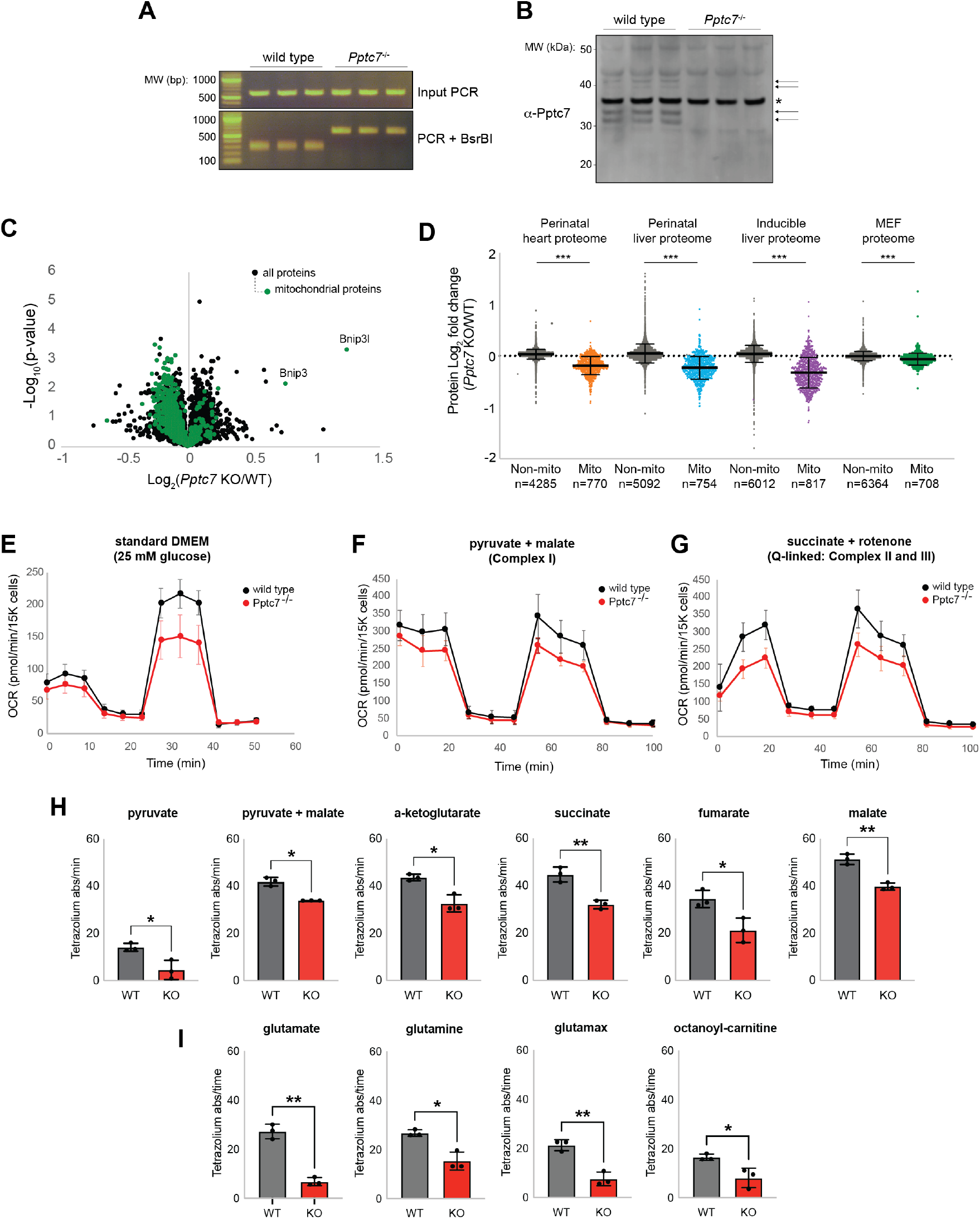
Pptc7 knockout causes cell-autonomous decreases in mitochondrial protein content leading to broad metabolic defects. **A**. Genotyping of wild type (WT) and *Pptc7* KO mouse embryonic fibroblasts (MEFs). **B**. Pptc7 endogenous protein expression in WT and KO MEFs. Notably, Pptc7 runs as a set of two doublets, marked by arrows. A non-specific band (*) serves as a loading control. **C**. Proteomic analysis of non-mitochondrial (black dots) and mitochondrial (green dots) proteins from triplicate WT and *Pptc7* KO MEFs. The mitophagy receptors Bnip3 and Nix are highlighted. **D**. Fold changes of mitochondrial proteins across all *Pptc7* KO systems including perinatal heart (orange) and liver (blue), inducible adult liver (purple), and MEFs (green). In each system, loss of *Pptc7* causes significant decreases in the mitochondrial proteome relative to non-mitochondrial proteins (shown in grey for all systems). ***p<0.001, one way ANOVA; multiple comparisons across each paired WT and KO dataset. Mean and standard deviation shown. **E**. Seahorse analysis of a mitochondrial stress test performed on primary *Pptc7* KO MEFs grown in standard DMEM supplemented with 25 mM glucose. **F**., **G**. Seahorse analysis of a mitochondrial stress test performed on primary permeabilized *Pptc7* KO MEFs given pyruvate and malate (**F**.) or succinate and rotenone (**G**). For all Seahorse analyses, error bars represent standard deviation. **H**.,**I**. BioLog analysis of permeabilized wild type and *Pptc7* KO MEFs given various TCA cycle substrates (H.) or amino acids and fatty acids (**I**.). For BioLog analysis, each datapoint represents an independent well on the BioLog plate, with the mean shown. Error bars represent standard deviation. *p<0.05, **p<0.01, ***p<0.001; BioLog data was analyzed using a Student’s t test.

We performed correlation analysis of all overlapping proteins identified in MEFs, perinatal heart tissue, and perinatal liver tissue, and found significant positive correlation between *Pptc7* KO models (Supplementary Figures 2A-B). These data also reveal that proteins involved in metabolic pathways, such as oxidative phosphorylation, the TCA cycle, and fatty acid oxidation were amongst the most decreased proteins across *Pptc7* KO cells and tissues (Supplementary Figures 2C-E), particularly when compared to other non-metabolic pathways (e.g., the mitochondrial ribosome) (Supplementary Figures 2F-H). To test whether *Pptc7* KO cells have compromised metabolism, we performed Seahorse assays and found that primary *Pptc7* KO fibroblasts have mildly decreased oxygen consumption rates (OCR) in basal conditions but have substantially lower spare respiratory capacity than wild-type fibroblasts (Figure 2E). We also tested OCR in permeabilized wild-type and *Pptc7* KO fibroblasts provided with pyruvate/malate or succinate/rotenone and found that KO cells given either substrate treatment showed decreased basal OCR and spare respiratory capacity (Figures 2F-G). Finally, we performed a BioLog assay to test the capacity of *Pptc7* KO cells to catabolize various nutrients. These experiments showed that *Pptc7* KO cells have a compromised ability to metabolize substrates that feed into the TCA cycle (e.g., pyruvate, α-ketoglutarate, succinate, fumarate, or malate, Figure 2H), glutamine catabolism (e.g., glutamate, glutamine, and Glutamax, Figure 2I), and fatty acid catabolism (e.g., octanoyl-carnitine, Figure 2I). Collectively, these data demonstrate that Pptc7 expression is required to maintain mitochondrial protein levels in isolated cells and that loss of this phosphatase disrupts metabolic function.

### Loss of mitochondrial content is mediated by the mitophagy receptors Bnip3 and Nix

The consistent loss of mitochondrial protein levels across each *Pptc7* KO tissue and cell type (Figure 2D) suggests a common molecular driver for this phenotype. The mitophagy receptor Bnip3 is significantly upregulated across all cell types and tissues assayed, with some systems, including the MEFs, showing additional upregulation of Nix (gene name *Bnip3l*), a Bnip3 paralog (Figures 1C, 2D, (Niemi et al., 2019)). We hypothesized that Bnip3 and Nix could promote excessive mitophagy, contributing to the decrease in mitochondrial protein levels seen in *Pptc7* KO systems. To test this, we used mt-Keima, a pH-dependent fluorescent protein that exhibits distinct emission spectra in mitochondria at physiological pH versus those present in the acidic lysosomal compartment after mitophagy (Sun et al., 2017). Indeed, *Pptc7* KO cells expressing mt-Keima showed elevated numbers of acidic mitochondria indicating excess mitophagic flux (Figures 3B-C). Importantly, this increase was fully abolished following CRISPR/Cas9-based disruption of *Bnip3* and *Bnip3l* (which can have functional redundancy in select contexts ((Zhao et al., 2020), Figures 3B-C) in the *Pptc7* KO background. Consistently, proteomic analysis of each of the *Pptc7/Bnip3/Bnip3l* knockout clones (herein referred to as TKO clones) revealed substantial rescue of most mitochondrial proteins relative to *Pptc7* KO cells (Figures 3D,E).

**Figure 3:**
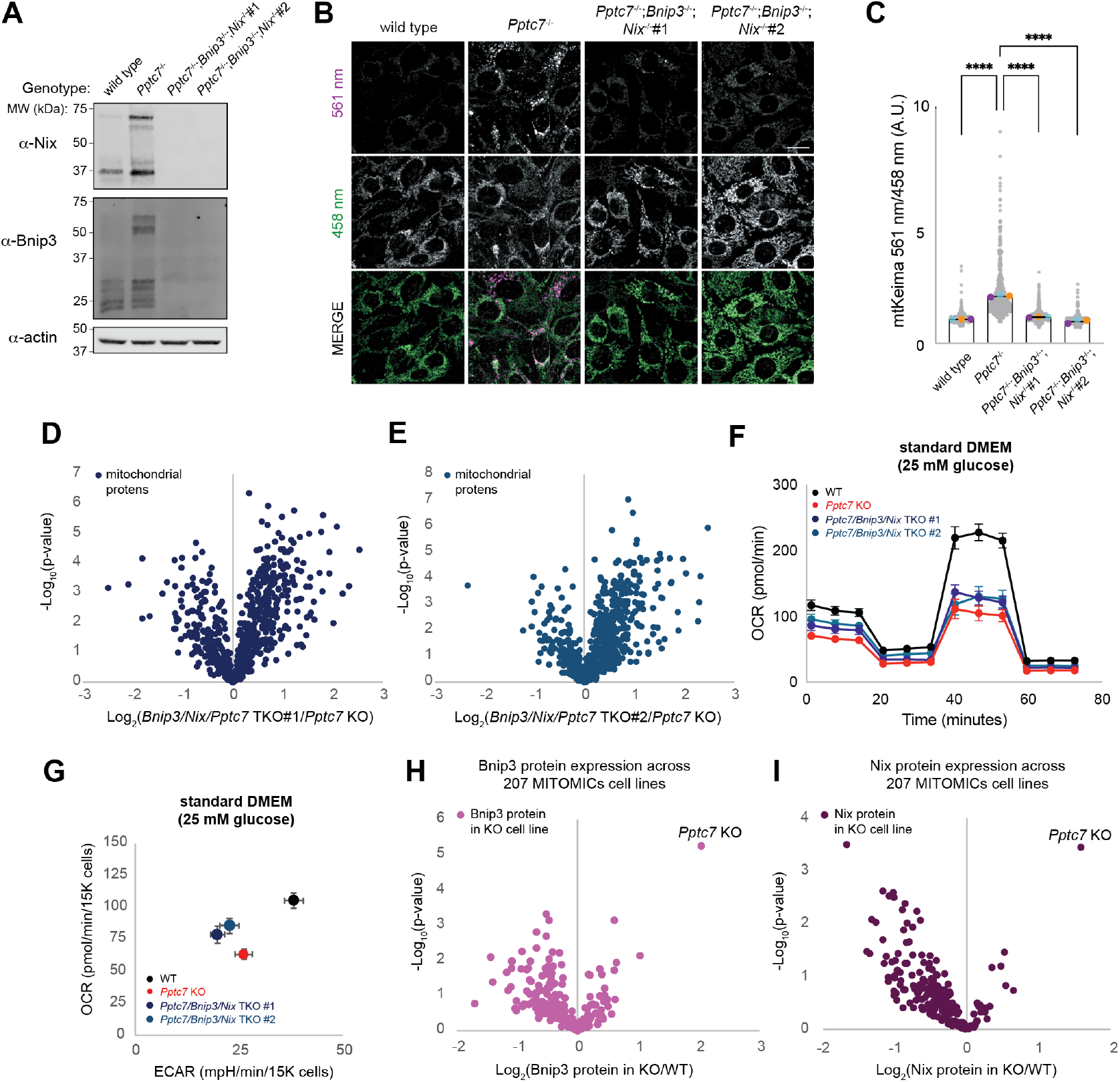
Pptc7 knockout causes excessive Bnip3- and Nix-mediated mitophagy. **A**. Western blot of Bnip3 and Nix expression in wild type, *Pptc7* KO, and two *Pptc7/Bnip3/Bnip3l* triple knockout (TKO) MEF cell lines. Actin is shown as a load control. **B**. Representative images of wild type, *Pptc7* KO, and each TKO cell line expressing the mitophagy reporter mt-Keima. Mitochondria imaged at 458 nm are at physiological pH but those imaged at 561 nm reflect acidic mitochondria undergoing mitophagy. **C**. Quantification of mt-Keima imaging. Each grey dot represents the ratio of mitochondrial fluorescence from a single cell. Blue, orange, and purple dots represent averages from three independent biological experiments. Error bars represent standard deviation. ****p<0.0001. mt-Keima data analyzed by one way ANOVA. **D**., **E**. Proteomic analysis of mitochondrial proteins in two independent *Pptc7/Bnip3/Bnip3l* TKO lines normalized to *Pptc7* KO alone. **F**. Seahorse analysis of a mitochondrial stress test in immortalized wild type (WT, blue), *Pptc7* KO (red), TKO #1 (dark blue), TKO #2 (teal). Cells were assayed in Seahorse DMEM supplemented with 25 mM glucose. Error bars represent standard deviation. **G**. Seahorse analysis of basal oxygen consumption rates (OCR) and extracellular acidification (ECAR). Error bars represent standard deviation. **H**., **I**. Bnip3 (**H**.) and Nix (**I**.) protein expression across 207 cell lines harboring monogenic mutations in genes encoding mitochondrial-localized proteins. Only *Pptc7* KO increases both Bnip3 and Nix significantly across this dataset.

We next tested whether the knockout of *Bnip3* and *Bnip3l* rescued the metabolic defects seen in *Pptc7* KO cells. Basal oxygen consumption was partially rescued in both TKO clones, but spare respiratory capacity was not rescued in either TKO clone relative to *Pptc7* KO cells (Figure 3F). Furthermore, the TKO cells exhibited diminished OCR and ECAR relative to wild-type MEFs (Figure 3G), similar to the *Pptc7* KO cells. These data demonstrate that while the dual knockout of Bnip3 and Nix reversed mitochondrial protein loss in the *Pptc7* KO cells, it did not rectify the underlying metabolic defects.

Due to the persistent metabolic defects seen in *Pptc7/Bnip3/Bnip3l* TKO cells, we considered the possibility that stabilization of Bnip3 and Nix may be an indirect, common downstream consequence of general mitochondrial dysfunction. To assess this, we analyzed the data from our recent MITOMICS study, in which we performed multiomic analyses on over 200 HAP1 cell lines carrying monogenic deletions of genes encoding mitochondria-localized proteins. Remarkably, *Pptc7* was the only gene whose KO resulted in significant increase of both Bnip3 and Nix expression in these cells (Figures 3H, I). These data suggest that upregulation of Bnip3 and Nix is not a general response to mitochondrial stress (at least in HAP1 cells), but rather that these mitophagy receptors are selectively elevated in response to the loss of *Pptc7*.

### Phosphoproteomic analyses reveal Bnip3 and Nix as candidate Pptc7 substrates

Our analyses above suggest that the metabolic aberrations caused by Pptc7 loss are rooted in dysregulated protein phosphorylation that then leads to Bnip3/Nix-mediated mitophagy. Our new model systems offer an opportunity to identify common Pptc7 substrates across cells and tissues. To identify these putative substrates, we performed phosphoproteomic analyses on liver tissue from both sexes of mice from our inducible *Pptc7* KO model (n=16 mice; 4 per sex and genotype, Figure 4A) and matched wild-type and *Pptc7* KO MEFs (n=3 independent cell lines for wild-type and KO, respectively, Figure 4B). As expected from the disruption of a protein phosphatase, the majority of significantly altered mitochondrial phosphoisoforms were elevated in *Pptc7* KO samples across both systems (Figures 4A and 4B). The results, especially when combined with our earlier data (Niemi et al., 2019), suggest a complex regulatory picture. Collectively, hundreds of mitochondrial phosphoproteins are elevated across all systems, including marked changes on multiple metabolic proteins. However, only four phosphoisoforms are significantly elevated across all systems (Figure 4C). Of note, there is substantially higher overlap of quantified phosphoisoforms in perinatal and adult model liver tissue from *Pptc7* KO mice that between other samples (Figure 4D), suggesting tissue specificity for Pptc7 substrates. These data are consistent with a recent study that found significant functional specialization of mitochondrial phosphoproteomes across tissue in mice (Mann et al., 2022). Validation of these potential substrates and their contribution to the *Pptc7* KO metabolic phenotype across tissues will require further mechanistic investigations.

**Figure 4:**
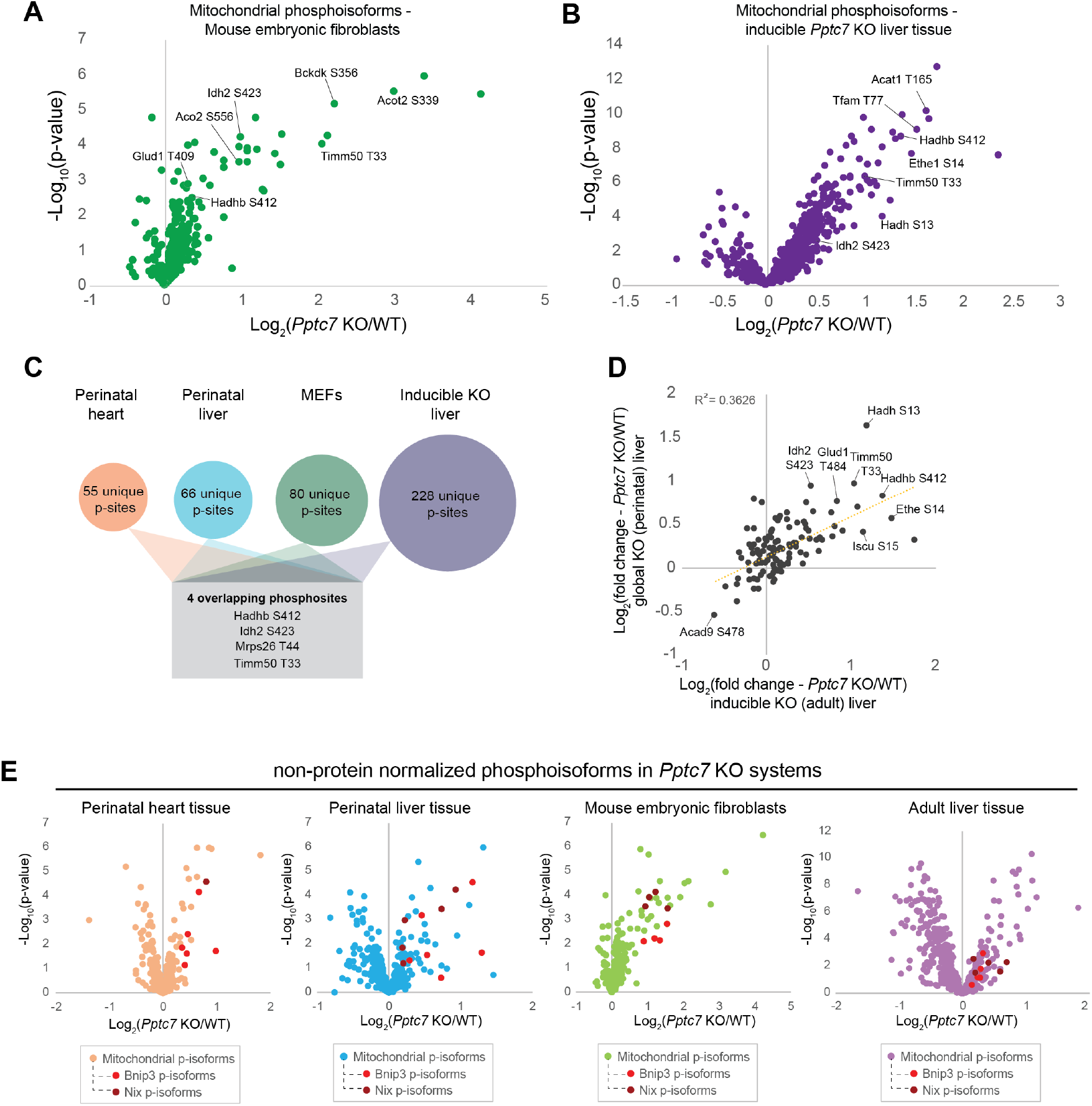
Phosphoproteomic analysis of Pptc7 KO systems reveals candidate substrates, including Bnip3 and Nix. **A**., **B**. Protein-normalized mitochondrial phosphoisoforms in *Pptc7* KO MEFs (green, **A**.) and inducible adult liver tissue (purple, **B**.). **C**. Analysis of phosphoproteomes across systems reveals many unique (i.e., only identified in a single experimental system) phosphosites (p-sites), with four significantly upregulated phosphosites identified across all four experimental systems (shown in grey box). **D**. Analysis of overlapping phosphosites identified in perinatal liver tissue (y-axis) and inducible adult liver (x-axis) show strong correlation. **E**. Non-protein normalized mitochondrial phosphoisoforms show Bnip3 and Nix have significantly elevated phosphorylation events across all tested model systems.

Notably, we identified multiple phosphorylation sites on the Bnip3 and Nix receptors themselves. These phosphorylation sites do not exhibit significant increases in the *Pptc7* KO cells when the data are subject to the standard normalization to total protein levels. However, this normalization could mask meaningful hits in instances whereby the phosphosite in question promotes protein stability and is found at high stoichiometry. In this case, loss of a phosphatase could result in the accumulation of stabilized, phosphorylated substrates that is not evident by a change in apparent phosphorylation occupancy. Given this, and the substantial upregulation of Bnip3 and Nix in our systems, we examined our phosphoproteomics data without total protein normalization, and found multiple elevated phosphopeptides for both Bnip3 and Nix across all *Pptc7* KO systems (Figure 4E). These phosphorylation events span eight unique residues in each of the Bnip3 and Nix proteins, suggesting complex phosphorylation-based regulation. Two of these observed sites have recently been associated with Bnip3 stability (He et al., 2022), adding further credence to the possibility that these phosphoisoforms may largely be representative of the total cellular pool of these proteins. Many aspects of this regulation remain to be elucidated, including the activity of kinase(s) responsible for Bnip3 and Nix phosphorylation, the effect of phosphorylation on protein half-life, and the absolute phosphorylation stoichiometry. Nonetheless, the consistent upregulation of these phosphopeptides in *Pptc7* KO systems suggests that Pptc7 influences this phosphorylation status, possibly through direct interaction.

## Discussion

Our data demonstrate that *Pptc7* knockout leads to Bnip3 (and, in some systems, Nix) upregulation, excessive mitophagy, and decreased steady state levels of mitochondrial proteins. Bnip3 and Nix are also phosphorylated in *Pptc7* knockout systems, consistent with a model in which phosphorylation promotes Bnip3- and Nix-mediated mitophagic signaling. Notably, in *S. cerevisiae*, the single mitophagy receptor Atg32 is phosphorylated by casein kinase 2 (CK2) to promote its interaction with Atg11 and subsequently induce mitophagy (Aoki et al., 2011; Kanki et al., 2013). Similar mechanisms activate mammalian mitophagy receptors, including FUNDC1, which is phosphorylated by CK2 and dephosphorylated through the phosphatase PGAM5 (Chen et al., 2014) and the Atg32 homolog Bcl2l13, whose phosphorylation allows recruitment of the mitophagy machinery (Murakawa et al., 2015). Our data suggest similar activation mechanisms may exist for Bnip3 and Nix, albeit more complex and potentially mediated through diverse signaling inputs, as evidenced by the numerous phosphorylation events identified on each receptor. Importantly, phosphorylation of Bnip3 and Nix has been previously linked to the function and stability of these proteins. Similar to FUNDC1, phosphorylation near the N-terminal LC3 interaction region (LIR) of Bnip3 and Nix stabilizes their interaction with LC3 to promote mitophagy (Poole et al., 2021; Rogov et al., 2017; Zhu et al., 2013). Phosphorylation has also been identified within the C-terminal regions of Bnip3 and Nix, influencing the ability of Bnip3 to promote cell death (Liu & Frazier, 2015) and the subcellular localization of Nix (da Silva Rosa et al., 2021). Finally, a recent study found that phosphorylation of Bnip3 at S60 and T66–two residues identified in our analysis–stabilizes this receptor by blocking its ubiquitination-mediated turnover (He et al., 2022). Notably, three independent studies recently identified the mitochondrial E3 ligase Fbxl4 as responsible for the basal turnover of Bnip3 and Nix (Elcocks et al., 2022; Jiang & Cao, 2022; Nguyen-Dien et al., 2022). Knockout of *Fbxl4* increases Bnip3 and Nix protein levels and promotes excessive mitophagy, leading to diminished mitochondrial protein levels and perinatal lethality in mice (Alsina et al., 2020)–phenotypes that strikingly mirror those observed in our *Pptc7* knockout models (Niemi et al., 2019).

Accumulating evidence suggests that cells devote significant resources to the proper regulation of Bnip3 and Nix activity. TMEM11, a transmembrane protein previously linked to MICOS complex, was recently found to physically interact with Bnip3 and Nix and mediate their spatial localization (Gok et al., 2023). Importantly, knockout of TMEM11 triggers hyperactive mitophagy, likely due to an increased number of LC3-positive mitophagosome formation sites at the outer mitochondrial membrane (Gok et al., 2023). These data suggest that the physical localization of Bnip3 and Nix is likely important for the proper regulation of mitophagy. Beyond this, Bnip3 protein levels seem to be actively limited in basal cell culture conditions, as Bnip3 exhibits one of the shortest-lived mitochondrial proteins with a half-life of 103 minutes (Schäfer et al., 2022). Bnip3 accumulates during proteasomal inhibition (He et al., 2022; Poole et al., 2021), suggesting efficient ubiquitin-mediated turnover. Consistently, *Fbxl4* knockout systems show hyperactive basal mitophagy, further suggesting that efficient ubiquitination of Bnip3 and Nix is critical for maintaining mitochondrial content and proper organellar function (Elcocks et al., 2022; Jiang & Cao, 2022; Nguyen-Dien et al., 2022). Our data, as well as data generated by He et al., suggest that phosphorylation of Bnip3 (and likely Nix) may promote protein stability by blocking ubiquitination-mediated turnover. However, He et al. identified PP1/2A as the phosphatase responsible for dephosphorylating Bnip3 (He et al., 2022). This leads to an important question: is Pptc7 directly responsible for the dephosphorylation of Bnip3 and Nix, or does it mediate these effects indirectly (for instance, through PP1/2A activity)?

Certain observations support each of these models. First, the matrix localization of Pptc7 (Rhee et al., 2013) should theoretically disallow the direct regulation of Bnip3 and Nix, which reside on the outer mitochondrial membrane (Hung et al., 2017; Rath et al., 2021). However, a handful of observations suggest that the location and function of Pptc7 may be more complex than currently appreciated. First, our western blot in Figure 2B reveals two sets of Pptc7 bands, one that migrates at the expected molecular weight (∼33 kDa), and one that migrates substantially higher than this (∼40 kDa). These data suggest that Pptc7 may reside in distinct populations in cells, although the molecular details underlying such differences in migration patterns remain to be elucidated. Second, the yeast ortholog of *Pptc7, PTC7*, is alternatively spliced to encode both a spliced protein that localizes to the mitochondrial matrix or a full-length protein that localizes to the ER or nuclear envelope (Juneau et al., 2009; Williams et al., 2014). These data suggest that Ptc7p and Pptc7 may share evolutionarily conserved dual functions in distinct organellar or cellular locations. Third, we recently found a physical interaction between PPTC7 and BNIP3 or NIX in a large-scale affinity enrichment-mass spectrometry analysis using mitochondrial baits in two human cell lines (Floyd et al., 2016). Our data revealed that the BNIP3 and NIX constitute the strongest identified interaction partners for PPTC7 in both 293T and HepG2 cells. Notably, of the 78 baits tested, PPTC7 is the only bait to pull down endogenous BNIP3 or NIX beyond our significance thresholds, suggesting this interaction is both specific and reproducible across experiments and cell types. These data are further supported by the BioPlex interactome studies (Huttlin et al., 2015, 2017, 2021), which also identify an interaction between PPTC7 and BNIP3. Notably, the BioPlex datasets also identify multiple interaction partners for PPTC7 in the mitochondrial matrix (e.g., DBT, RTN4IP1, TRMT2B, and TRMT61B) as well as on the outer mitochondrial membrane (e.g., RHOT1, RHOT2, HSDL1, MAOB, and TRABD), which may support dual localization for this phosphatase. We propose that these data, along with the data presented in this manuscript, support a role for Pptc7 in the direct dephosphorylation of Bnip3 and Nix. Which residues on Bnip3 and Nix are dephosphorylated, whether these interactions are direct, and the cellular contexts in which these interactions take place should be an active area of investigation in the future.

Despite the importance of Bnip3 and Nix in mediating excessive mitophagy in *Pptc7* knockout systems, the lack of full metabolic rescue in our TKO cells suggests that this phosphatase influences mitochondrial functions in ways distinct from its functional interaction with these mitophagy receptors. Consistently, Bnip3 and Nix are not conserved in *S. cerevisiae*, while Pptc7 has a functional yeast ortholog, PTC7, that promotes mitochondrial function (Guo et al., 2017; Martín-Montalvo et al., 2013). This suggests that Pptc7 has ancestral roles in enabling mitochondrial metabolism independent of its role in mammalian mitophagy. One such function may be the regulation of mitochondrial protein import via Timm50. Phosphorylation at threonine 33 of Timm50 is one of the four consistently upregulated phosphorylation events in *Pptc7* KO systems (Figure 4C), and phosphorylation in a similar region of Tim50p (at serine 104) was identified in *ptc7*Δ yeast (Guo et al., 2017; Niemi et al., 2019). We previously demonstrated that phosphomimetic mutation of S104 in Tim50p decreases mitochondrial protein import and that a non-phosphorylatable S104A Tim50p mutant partially rescues import defects in *ptc7*Δ yeast (Niemi et al., 2019). These data suggest that the dysregulated phosphorylation on Timm50, or other candidate substrates identified in our analyses, likely contributes to the metabolic dysfunction seen in *Pptc7* KO systems. The TKO cells generated for this study will serve as a critical tool for disentangling these metabolic phenotypes in the absence of diminished mitochondrial protein content.

Collectively, our data suggest that loss of the mitochondrial phosphatase Pptc7 decreases mitochondrial protein content in isolated cells, in multiple tissue types, and across developmental contexts. This loss in mitochondrial protein content is associated with elevated phosphorylation on numerous mitochondrial proteins, including the mitophagy receptors Bnip3 and Nix. The decreases in mitochondrial protein content seen in *Pptc7* KO cells are largely rescued by genetic ablation of Bnip3 and Nix without rectifying the underlying metabolic issues. These data highlight the critical role of Pptc7 in maintaining mitochondrial homeostasis, and further substantiate the importance of properly regulated protein phosphorylation within this organelle.

## Methods

### Generation of a conditional floxed Pptc7 mouse model

*Pptc7* floxed animals were generated at the Biotechnology Center at the University of Wisconsin-Madison. The *Easi*-CRISPR method (Quadros et al., 2017) was used to generate two loxP sites flanking exon 3 of Pptc7 (i.e., a floxed allele). Animals were generated with a CRISPR approach as previously described (Niemi et al. 2019), with a key modification the addition of a long, single stranded oligonucleotide (i.e., a megamer) to facilitate homologous recombination. Briefly, one-cell fertilized C57BL/6J embryos were microinjected with CRISPR reagents (ctRNA.int12.2 (20ng/ul) + ctRNA.int23.3 (20ng/ul) + Cas9 IDT (50ng/ul) + megamer (50ng/ul)). Sequences of ctRNA.int12.2, ctRNA.int23.3 and the sequence of the resulting floxed allele is provided in Supplementary Table 1. Microinjected embryos were surgically transferred into pseudeopregnant recipients. Pups were sequenced at weaning for each LoxP cassette. Founders were backcrossed to C57BL/6J mates and F1 pups analyzed by sequencing to evaluate germline transmission.

### Generation of an inducible global Pptc7 knockout mouse

*Pptc7* floxed animals were bred with UBC-Cre-ERT2 transgenic mice to generate animals a global, inducible Pptc7 knockout model. The UBC-Cre-ERT2 strain was acquired from Jackson laboratories (B6.Cg-*Ndor1*^*Tg(UBC-cre/ERT2)1Ejb*^/1J, Strain #007001) and was bred to Pptc7^flox/flox^ animals. Pptc7 floxed animals were bred to homozygosity (i.e., Pptc7^flox/flox^) while maintaining one copy of the UBC-Cre/ERT2 transgene (referred to as ‘Cre’ hereafter) as experimental animals. Control animals include UBC-Cre/ERT2;Pptc7^+/+^ animals and/or Pptc7^flox/flox^ animals carrying no copy of Cre transgene. Controls used in each experiment are clarified within the figure legends of the manuscript. Both male and female mice were used in our analyses. To induce knockout, mice were fed tamoxifen-containing chow for seven consecutive days. Tamoxifen chow was purchased from Envigo (Teklad custom diet TD.130859), which is formulated for 400 mg. of tamoxifen citrate per kilogram of diet. The diet was modified to promote more consistent feeding and genomic knockout as follows: tamoxifen chow pellets were ground to a powder with a morter and pestle, as was standard chow (Formulab Diet 5008). The two diets were mixed 3:1 (i.e., 15 g. of tamoxifen chow to 5 g. of standard chow) and placed into a weigh boat. Water was added to the powder to create a “mash”; a fresh mash was given to each cage daily for the duration of the treatment. After tamoxifen administration was complete, mice were transitioned back to a standard chow diet (Formulab Diet 5008) until the end of the experiment.

### Genotyping analysis

Each model used in our studied was validated by genotyping analysis. Genotyping primers for the flox allele and Cre used for genotyping can be found in Supplementary Table 2. The flox primers produce a 667 bp amplicon for the floxed allele and a 599 bp amplicon for the wild type allele. Upon Cre-mediated excision, these primers produce a 167 bp amplicon reflecting a knockout allele. The Cre-specific primers produce a 480 bp amplicon for the Cre allele. Approximately 100 ng of genomic DNA isolated from tail tips served as a template for all genotyping reactions. *Pptc7* knockout MEFs were generated from our previously published CRISPR knockout model and genotyped as previously described (Niemi et al., 2019).

### Generation and validation of mouse embryonic fibroblasts

Mouse embryonic fibroblasts were generated from a cross of Pptc7^+/-^ heterozygous mice. A timed mating was performed and E0.5 was determined through the presence of a vaginal plug. The pregnant female was sacrificed at E14.5 by CO_2_ asphyxiation. 11 embryos were removed, decapitated, and internal organs removed. Remaining tissue was finely minced, suspended in 0.25% trypsin-EDTA, and incubated at 37°C for 30 minutes in a CO_2_-controlled tissue culture incubator. After trypsinization, tissues were pipetted to a single cell suspension, filtered through a nylon mesh cell strainer (Thermo-Fisher #08-771-1), and plated into individual 60 mm^2^ dishes. Cells were cultured in DMEM (high glucose, no pyruvate, Thermo-Fisher catalog #11965-092) supplemented with 10% heat inactivated fetal bovine serum (FBS) and 1x penicillin/streptomycin. Cells were grown in a temperature-controlled CO_2_ incubator at 37°C and 5% CO_2_. For the first 3 passages, media was supplemented with gentamicin (10 μg/ml) to reduce risks of contamination. Each cell line was genotyped using leftover tissue from each embryo from which genomic DNA was isolated. gDNA isolation and genotyping was done as previously described (Niemi et al., 2019). The sex of the cells was not determined as male/female embryos are morphologically indistinguishable at E14.5. Immortalized MEFs were generated using plasmid encoding SV40 Large T antigen as follows: MEFs were split into 6 well plates to ∼80% confluence and transfected with 2 µg SV40 1: pBSSVD2005 (a gift from David Ron; Addgene plasmid #21826) using FuGENE HD according to manufacturer’s protocol. MEFs were split 1:10 for 5 consecutive passages before considered immortalized. Immortalized MEFs were cultured identically to primary MEFs (see above) and were used to a maximum of 15 passages for all experiments. Cellular experiments in this paper were performed with both primary (e.g., Seahorse assays) and immortalized (e.g., phosphoproteomics, proteomics, and BioLog assays) MEF clones.

### Proteomic and phosphoproteomic analysis

TMT-based proteomic and phosphorproteomic experiments were performed on mouse liver tissue and mouse embryonic fibroblasts, Supplemental datasets 1 and 2, respectively. Label-free proteomics analysis was performed on *Pptc7/Bnip3/Bnip3l* TKO samples, Supplemental dataset 3. Details for the generation of these datasets are as follows:

### TMT phosphoproteomics

#### Tissue lysis and protein digestion

Cells or mouse liver tissue were resuspended in 1 mL 6 M guanidine hydrochloride, 100 mM Tris, pH 8 and probe-sonicated using a Misonix XL-2000 Series Ultrasonic Liquid Processor. The sonication regime, repeated twice, was the following: the probe was set to output 11 watts for 10 seconds with an off-time of 30 seconds. The protein concentrations of the homogenized tissue samples were determined using the BCA Protein Assay Kit (Thermo Pierce). Sample volumes corresponding to 400 μg of protein were set aside for further processing. Methanol was then added to each sample so that the final volume consisted of 90% methanol. The samples were then centrifuged at 9,000 g for 5 minutes, and the supernatant was decanted. The protein pellet was then resuspended in 200 μL of lysis buffer containing 8 M urea, 10 mM tris (2-carboxyethyl) phosphine, 40 mM chloroacetamide, and 100 mM Tris, pH 8. 8 μg of Lysyl Endopeptidase (FUJIFILM Wako Chemicals) was added and incubated for 4 hours at room temperature. The samples were then diluted with 100 mM tris, pH 8 to achieve a final urea concentration of less than 2 M. 8 μg of trypsin (Promega) was added, and the samples incubated for 10 hours at room temperature. Trypsin enzyme activity was then quenched by addition of 120 μL of 10% trifluoroacetic acid (TFA), and the samples were centrifuged at 9,000 g for 5 minutes. Peptides were desalted with Strata-X reversed phase solid phase extraction cartridges (Phenomenex).

#### TMT labeling

Desalted peptides were resuspended in 100 μL 200 mM triethylammonium bicarbonate buffer. Each TMTpro 16 Plex channel had been pre-aliquoted in 500 μg quantities was resuspended in 20 μL of acetonitrile (ACN). Each sample was added to distinct TMT channels and vortexed for 4 hours at room temperature. The TMT labeling reaction was quenched with 0.64 μL of 50% hydroxylamine and 100 μg of labeled peptides per sample were combined. The combined sample was dried and desalted with Strata-X reversed phase solid phase extraction cartridges.

#### Phosphopeptide enrichment

Desalted TMT-labeled peptides were resuspended in 1 mL 8% can in 6% TFA. 104 μL of magnetic titanium dioxide beads (ReSyn Biosciences) that were washed with 80% ACN 6% TFA three times. The resuspended sample was added to the washed beads and vortexed for 1 hour at room temperature. The bead-bound phosphorylated peptides were then sequentially washed three times with 1 mL of 80% ACN/6% TFA, once with 80% ACN, once with 80% ACN in 0.5M glycolic acid, then three times with 80% ACN. The phosphorylated peptides were eluted from the beads with two washes of 300 μL 50% ACN in 1% ammonium hydroxide. Dried phosphorylated peptides were resuspended in 0.1% TFA and desalted using Strata-X reversed phase solid phase extraction cartridges.

#### Offline fractionation

The TMT-labeled phosphopeptides were loaded onto a 4.6 mm inner diameter 150 mm outer diameter BEH C18 column (Waters Corp) for offline fractionation into 16 fractions. The peptides were separated using a high pH reversed-phase gradient (10 mM ammonium formate Mobile Phase A, 10 mM ammonium formate in 80% MeOH Mobile Phase B) on an Agilent 1260 Infinity Binary LC equipped with an automatic fraction collector. The effective separation gradient started at 20% Mobile Phase B and increased to 60% Mobile Phase B in 6 minutes, flowing at a rate of 800 μL/min. The fractions were then concatenated into 8 fractions, combining fractions 1 and 9, 2 and 10, 3 and 11, 4 and 12, 5 and 13, 6 and 14, 7 and 15, and 8 and 16. The combined fractions were transferred to 2 mL Starstedt micro tubes for rapid evaporation under vacuum with the SpeedVac Vacuum Concentrator Kit (Thermo).

#### LC-MS acquisition

Each fraction was resuspended in 0.2% formic acid. 1 μg of peptide was loaded onto a 75 μm inner diameter x 360 μm outer diameter column (New Objective) packed in-house 1 to 30 cm with 1.7 micron BEH C18 particles (Waters Corp). The peptides were chromatographically separated over a 120-minute reverse phase gradient (0.2% formic acid mobile phase A, 0.2% formic acid/80% ACN mobile phase B) using a Dionex UltiMate 3000 nano-HPLC (Thermo) at a column temperature of 50°C and a flow rate of 335 nL/min. Eluting peptides were sprayed into an Orbitrap Eclipse using a Nanospray Flex ionization source (Thermo) at 2 kV. MS1 scans were recorded in the Orbitrap using a resolving power of 60,000, requiring an AGC target of 1e6 or a maximum injection time of 50 ms. MS1 scans included precursor ions ranging from 300-1350 m/z. The most intense ions were selected for HCD fragmentation (normalized collision energy 35%) and subsequent MS2 analysis until the 1 second duty cycle lapsed. Dynamic exclusion was set to 40 seconds and the MS2 isolation width was set to 1.5 Th. Only precursors of charge states 2-6 were selected. MS2 spectra were analyzed in the Orbitrap using a resolving power of 60,000 requiring an AGC target of 1e5 or a maximum inject time of 118 ms. MS2 scans included fragment ions ranging from 100-2000 m/z.

#### Data Processing

Thermo RAW files were processed with MaxQuant (version 1.5.2.8)2, implementing the Andromeda3 algorithm to perform database searching of MS2 spectra against a database of canonical proteins and isoforms (Uniprot, Mus musculus, 2018). TMT labels were set as fixed modifications on N termini and lysine residues. Carbamidomethylation of cysteine was set as a fixed modification. Variable modifications included acetylation of the N terminus, phosphorylation of serine, threonine, and tyrosine, and oxidation of methionine. Peptide and protein matches were filtered to 1% FDR. Phosphorylation sites were considered localized if they delivered MaxQuant localization scores >0.75. The TMT reporter ion intensities of the phosphorylated peptides were median-normalized to account for variability introduced from sample-handling. To this end, the median of the reporter ion intensities for each sample was calculated. To obtain the correction factor for each phosphorylated peptide intensity, the average of the medians was divided by each of the respective sample medians, rendering a correction factor for each channel. This channel-specific correction factor was then multiplied to each of the reporter ion intensities within that channel.

### Label-free proteomics

#### Protein extraction and digestion

Mouse embryonic fibroblast cells were resuspended in 1 mL 6 M guanidine hydrochloride, 100 mM tris, pH 8 and sonicated using a Qsonica Q700 temperature-controlled sonicator. The sonication regime cycled 10 times applying an ultrasonic wave of amplitude of 35 for 20 seconds with a 10 second off time. The protein concentrations of the lysate were then determined using the Pierce BCA Protein Assay Kit. Methanol was then added to volumes corresponding to 200 μg for each sample, so that the final volume consisted of 90% methanol for protein extraction. The samples were then centrifuged at 9,000 g for 5 minutes and the supernatant was decanted. The protein pellet was then resuspended in 100 μL of lysis buffer containing 8M urea, 10 mM tris (2-carboxyethyl) phosphine, 40 mM chloroacetamide, and 100 mM tris, pH 8. 4 μg of Lysyl Endopeptidase was added and incubated for 4 hours at room temperature. The samples were then diluted with 100 mM tris, pH 8 to achieve a 2 M urea concentration. 4 μg of trypsin was added and the samples incubated for 10 hours at room temperature. Trypsin enzyme activity was then quenched by adding 60 μL of 10% TFA and the samples were centrifuged at 9,000 g for 5 minutes. Peptides were desalted with Strata-X reversed phase solid phase extraction cartridges.

#### LC-MS acquisition method

Each sample was resuspended in 0.2% formic acid. 1.5 μg of peptide was loaded onto a 75 um inner diameter x 360 um outer diameter column (CoAnn Technologies) packed in-house1 to 30 cm with 1.7 micron BEH C18 particles (Waters Corp). The peptides were chromatographically separated over a 90-minute reverse phase gradient (0.2% formic acid mobile phase A, 0.2% formic acid/80% ACN mobile phase B) using a Vanquish Neo UHPLC (Thermo) at a column temperature of 50°C and a flow rate of 320 nL/minute. Eluting peptides were sprayed into an Orbitrap Eclipse using a Nanospray Flex ionization source (Thermo) at 2.2 kV. MS1 scans were recorded in the Orbitrap using a resolving power of 240,000, requiring an AGC target of 8e5 or a maximum injection time of 50 ms. MS1 scans included precursor ions ranging from 350-2000 m/z. The most intense ions were selected for HCD fragmentation (normalized collision energy of 25%) and subsequent MS2 analysis until the 1 second duty cycle lapsed. Dynamic exclusion was set to 10 seconds and the quadrupole isolation width was set to 0.5 Th. Only precursor ions with charge states of 2-5 were selected. MS2 spectra were analyzed in the ion trap on the “turbo” resolution setting, requiring an AGC target of 2e4 or a maximum inject time of 14 ms. MS2 scans included fragment ions ranging from 150-1350 m/z.

#### Data Processing

Thermo RAW files were processed with MaxQuant (version 1.5.2.8)2, implementing the Andromeda3 algorithm to perform database searching of MS2 spectra against a database of canonical proteins and isoforms (Uniprot, Mus musculus). Carbamidomethylation was set as a fixed modification on cysteines. Variable modifications included N terminus acetylation and methionine oxidation. Peptide and protein matches were filtered to 1% FDR. Fast LFQ was toggled off and a LFQ minimum ratio count was set to 1. Match between runs (MBR) was toggled on. All other parameters were set to default MaxQuant settings.

#### Metabolic assays

For the Seahorse assays in Figure 2, primary (i.e., not immortalized) *Pptc7* knockout MEFs and matched wild-type MEFs were plated into Seahorse XFe96 plates at 12,500 cells/well. For the Seahorse assay in Figure 3, immortalized wild-type, *Pptc7* knockout, and *Pptc7/Bnip3/Bnip3l* triple knockout cells were plated into XFe96 plates at 15,000 cells/well. Cells were allowed to adhere overnight in standard DMEM media (25 mM glucose, 2 mM glutamine, 10% heat inactivated FBS, 1x penicillin/streptomycin). One hour before the Seahorse experiment, DMEM was aspirated, cells were washed with PBS, and were swapped into Seahorse XF Base Medium (i.e., unbuffered DMEM containing 25 mM glucose, 2 mM glutamine with no FBS) and allowed to equilibrate at 37°C in a non-CO_2_ incubator before initiating the experiment. While cells were equilibrating, a pre-hydrated Seahorse cartridge was loaded with oligomycin, FCCP, and Rotenone/antimycin A prepared from a XF Cell Mito Stress Test Kit, which were reconstituted per the manufacturer’s directions. Compounds were injected into 180 ul assay medium in subsequent injections for final concentrations of 1.0 μM oligomycin, 1.0 μM FCCP, and 0.5 μM rotenone/antimycin A. For pyruvate/malate and succinate/rotenone experiments, primary *Pptc7* knockout and wild type MEFs were permeabilized using XF PMP per manufacturer’s instructions. MEFs were plated in a 96 well plate as described above. Plating DMEM was aspirated, and cells were washed with 1x MAS (220 mM mannitol, 70 mM sucrose, 10 mM KH2PO4, 5 mM MgCl2, 2 mM HEPES, 1 mM EGTA, and 0.2% w/v fatty acid free BSA). To initiate the experiment, cells were swapped into 1x MAS containing 1 nM XF PMP (to permeabilize the cells) and either 1 mM malate and 10 mM pyruvate or 10 mM succinate and 2 μM rotenone. Permeabilized cells were subjected to the XF Cell Mito Stress Test Kit as described above. All data were collected on an XFe96 Seahorse Flux analyzer and analyzed on its associated Seahorse XF software. To perform the BioLog experiment, a BioLog MitoPlate S-1 Mitochondrial Substrate Metabolism assay plate was activated by adding ‘assay mix’ (BioLog MAS, Redox Dye MC, saponin, and water) to each well and incubating for 1 hour at 37°C to dissolve substrates. Immortalized *Pptc7* knockout MEFs and matched wild type MEFs were trypsinized, counted to a final cell number of 20,000 cells per well, and resuspended in 3 ml of 1x BioLog MAS (Biolog catalog #72303). The redox dye color formation (OD590) within the plate was tracked on an OmniLog instrument for 4 hours. Data were collected on OmniLog software and analyzed by calculating the slope of redox dye formation as reported over this 4-hour timeframe.

#### Citrate synthase assays

Citrate synthase assays were performed as previously described (Guo et al., 2017; Niemi et al., 2019). Briefly, liver tissue was homogenized in Lysis Buffer A (LBA; 50 mM Tris-HCl, pH 7.4, 40 mM NaCl, 1 mM EDTA, and 0.5% (v/v) Triton X-100) supplemented with 1x protease inhibitor cocktail (0.5 μg/ml pepstatin A, chymostatin, antipain, leupeptin, and aprotinin) and 1x phosphatase inhibitor cocktail (0.5 mM imidazole, 0.25 mM sodium fluoride, 0.3 mM sodium molybdate, 0.25 mM sodium orthovanadate, and 1 mM sodium tartrate). Lysates were clarified, BCA normalized, and 5 μg of whole cell lysate was used for each reaction. 200 μl reactions were assembled into a single well of a 96 well plate containing the following (final concentrations): 100 mM Tris, pH 7.4, 300 μM acetyl CoA, 100 μM DTNB (5,5’-dithio-bis-[2-nitrobenzoic acid]). 10 µL of 10 mM oxaloacetic acid (OAA) was added per well (final [c] = 500 μM) to initiate enzyme assay. Absorbance at 412 nm (A412) was measured every minute for 30 minutes on a Cytation 3 plate reader (BioTek).

#### Western blotting and antibodies

Western blotting was performed as previously described (Niemi et al., 2019). Antibodies used in this study include: anti-Pptc7 (Novus Biologicals, catalog #NBP1-90654, 1:1000 dilution, 48 hour incubation), anti-Citrate synthase (Cell Signaling Technologies, catalog #14309, 1:1000 dilution, overnight incubation), anti-beta actin (Abcam, catalog # ab170325, 1:1000 dilution, overnight incubation), anti-Bnip3 (rodent specific antibody, Cell Signaling Technologies, catalog #3769, 1:1000 dilution, 48 hour incubation), and anti-Nix (Cell Signaling Technologies, catalog #12396, 1:1000 dilution, overnight incubation).

#### mtDNA analysis

Genomic DNA (i.e., gDNA) was isolated from mouse liver tissue using the DNeasy Blood & Tissue Kit (Qiagen) according to manufacturer’s instructions. DNA was quantified and normalized to 100 ng input per reaction. Primers to analyze nuclear and mitochondrial DNA markers can be found in Supplemental Table 3. Quantitative real time PCR was performed on a QuantStudio 6 Real-Time PCR system with data was collected on QuantStudio Real-Time PCR v1.2 software. Relative quantitation was performed using the ΔΔCt method.

#### Generation of Pptc7/Bnip3/Nix triple knockout (TKO) cells

A single clone of *Pptc7* knockout MEFs were used to generate *Pptc7/Bnip3/Bnip3l* triple knockout (TKO) cells. The TKO MEFs were created by the Genome Engineering & Stem Cell Center (GESC@MGI) at Washington University in St. Louis. Briefly, two gRNAs were designed to target each 5’ and 3’ of exon of interest, introduce double strain break and delete target exon. Synthetic gRNAs were purchased from IDT (sequences are provided in Supplemental Table 4), complexed with Cas9 recombinant protein and transfected into MEFs. The transfected cells were then single cell sorted into 96-well plates. Single cell clones were identified using deletion PCR and inside PCR to analyze the target site region for complete deletion of the respective gene (primers sequence and PCR products size are reported in Supplemental Table 4).

#### Mitophagy assays with mt-Keima

The mitochondrial keima (mt-Keima) assay was described previously in (Sun et al., 2017). Live-cell images were acquired using a Leica DMi8 SP8 Inverted Confocal microscope, equipped with a 63x Plan Apochromatic objective and environmental chamber (5% CO2, 37°C). Images were quantified with Image J/FIJI software. Single cells were segregated by generating regions of interest (ROI). Selected ROIs were cropped and split into separated channels for 561 nm and 458 nm, followed by threshold processing. The fluorescence intensity of mt-Keima at 561 nm (lysosomal signal) and mt-Keima at 458 nm (mitochondrial signal) was measured for each ROI and the ratio (561 nm/458 nm) was calculated. Three biological replicates were performed, with >50 cells analyzed per condition for each replicate. Individual ratios are represented as grey dots and the mean ratios from each biological replicate.

## Acknowledgements

We would like to thank members of the Pagliarini lab and the Niemi lab for discussions and critical evaluation of this work. This work was supported by 5R01DK098672 (to D.J.P.), P41GM108538 (National Center for Quantitative Biology of Complex Systems, to D.J.P. and J.J.C.), 5T32G002760 (Genomic Sciences Training Program training grant support to L.R.S.), and funds from the BJC Investigators Program (to D.J.P.). Support for this research was provided by the University of Wisconsin–Madison, Department of Biochemistry and Office of the Vice Chancellor for Research and Graduate Education with funding from the Wisconsin Alumni Research Foundation (to M.P.K.), and by NIH grants, R01DK101573, R01DK102948, and RC2DK125961 (to A.D.A.). J.K.P. was supported by Australian National Health and Medical Research Council grants (APP1183915 and APP 2019993), a Brain Foundation Research grant (2020), and an Australian Research Council Future Fellowship (FT180100172). J.K.P and K.K. were supported by a Mito Foundation Incubator Grant (2022) and Mito Foundation Ph.D. scholarship, respectively. We would like to thank Kathy Krentz and C. Dustin Rubenstein from the Animal Models and Genome Editing Cores at the University of Wisconsin-Madison for generating of the *Pptc7* floxed model. The generation of the *Pptc7* floxed model was supported by the University of Wisconsin Carbone Cancer Center Support Grant (P30CA014520). We thank the Genome Engineering and iPSC Center (GESC@MGI) at the Washington University in St. Louis for creating the *Pptc7/Bnip3/Bnip3l* triple knockout (TKO) MEFs lines. We would like to acknowledge the Washington University Diabetes Research Center (P30DK020579) for providing access to the Seahorse instrument used to generate data in this manuscript.

## Conflict of interest

The following conflicts of interest have been declared: J.J.C. is a consultant for Thermo Fisher Scientific, 908 Devices, and Seer.

---

**Supplementary Figure 1:**
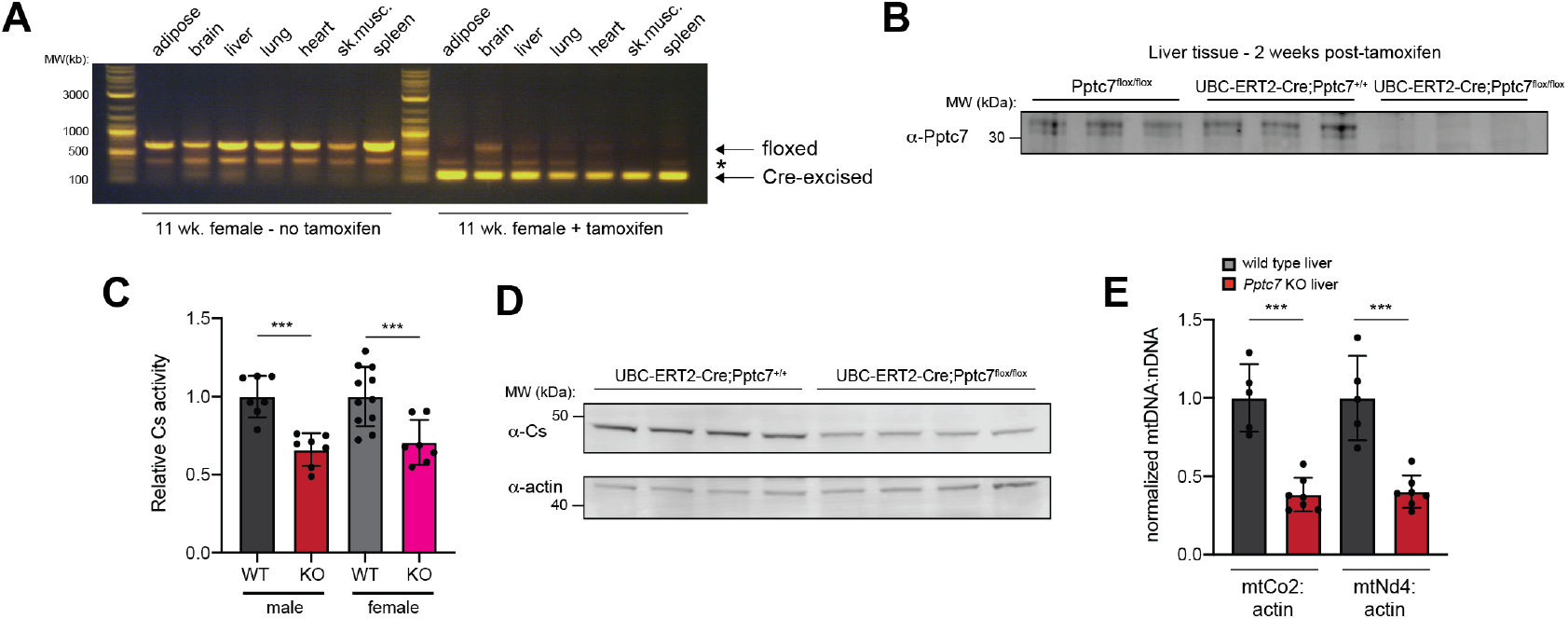
Establishment and validation of the inducible Pptc7 KO knockout mouse. **A**. Genotyping for the floxed allele in 11 week old female mice without (left) and with (right) one week of tamoxifen administration. Genotyping reveals substantial recombination only in the presence of tamoxifen for most tissues assayed. *represents a non-specific band. **B**. Endogenous Pptc7 in floxed mice (left, n=3), Cre-containing wild type mice (middle, n=3), or experimental knockout animals (right, n=3). Only the experimental animals show knockout at the protein level. **C**. Citrate synthase (Cs)) activity in liver tissue from male and female mice. Each dot represents Cs activity from an individual animal; error bars represent standard deviation. ***p<0.001, Student’s t test. **D**. Expression of citrate synthase protein in n=4 control (left) or experimental (right) animals. Actin is shown as a load control. **E**. Relative mtDNA levels (compared to nuclear DNA, or nDNA) in male wild type (grey) or *Pptc7* KO (red) liver. Error bars represent standard deviation. ***p<0.001, Student’s t test.

**Supplementary Figure 2:**
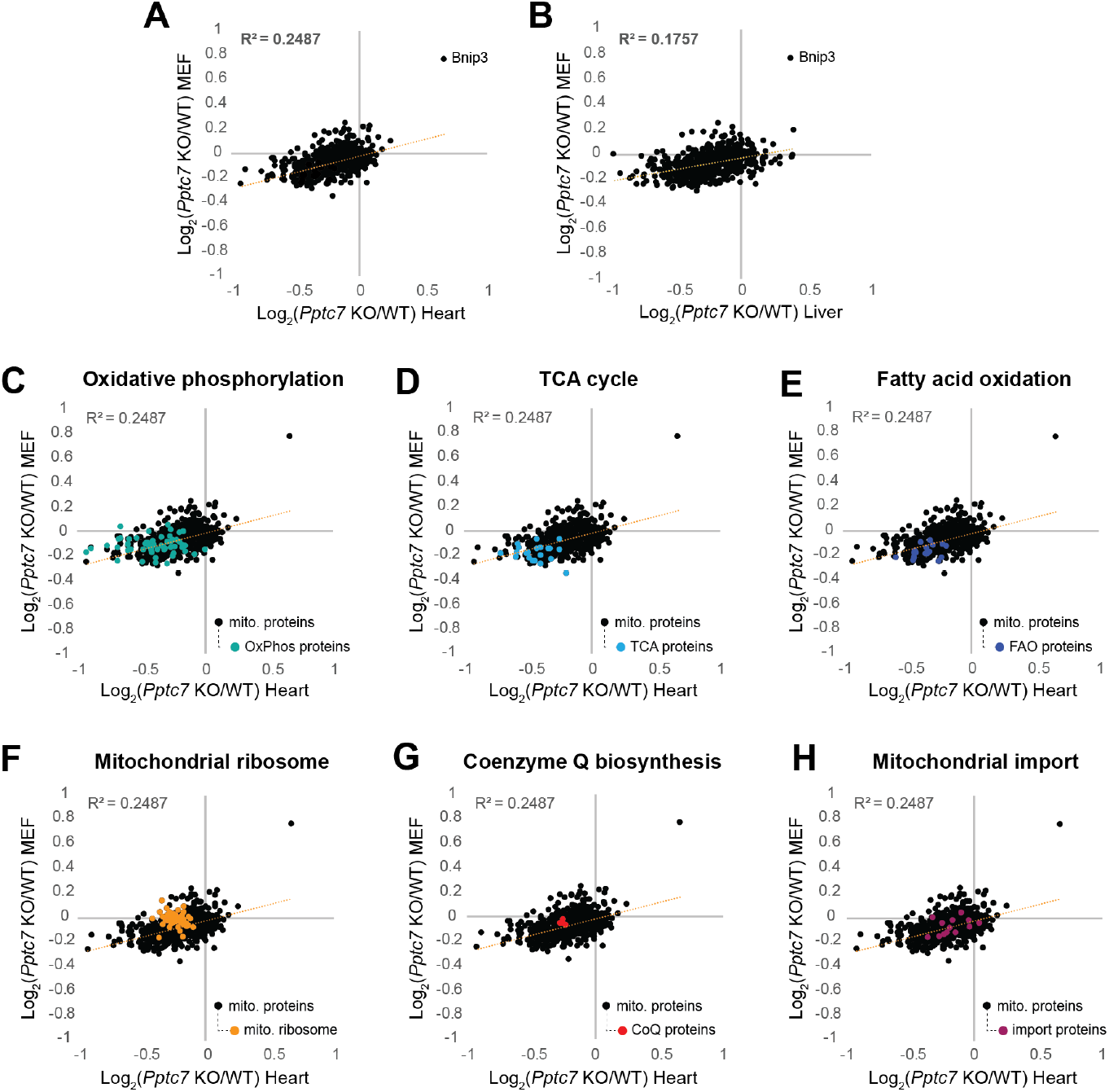
Metabolic proteins are amongst the most affected by Pptc7 KO. **A**., **B**. Correlation analysis between mitochondrial protein fold changes in *Pptc7* KO MEFs (y-axis) and perinatal heart (x-axis, **A**.) and perinatal liver (x-axis, **B**.). **C**.-**H**., Pathway analysis highlighting proteins involved in oxidative phosphorylation (**C**.), the TCA cycle (**D**.), fatty acid oxidation (**E**.), the mitochondrial ribosome (**F**.), coenzyme Q biosynthesis (**G**.), and mitochondrial protein import (**H**.).

**Supplementary Table 1.**
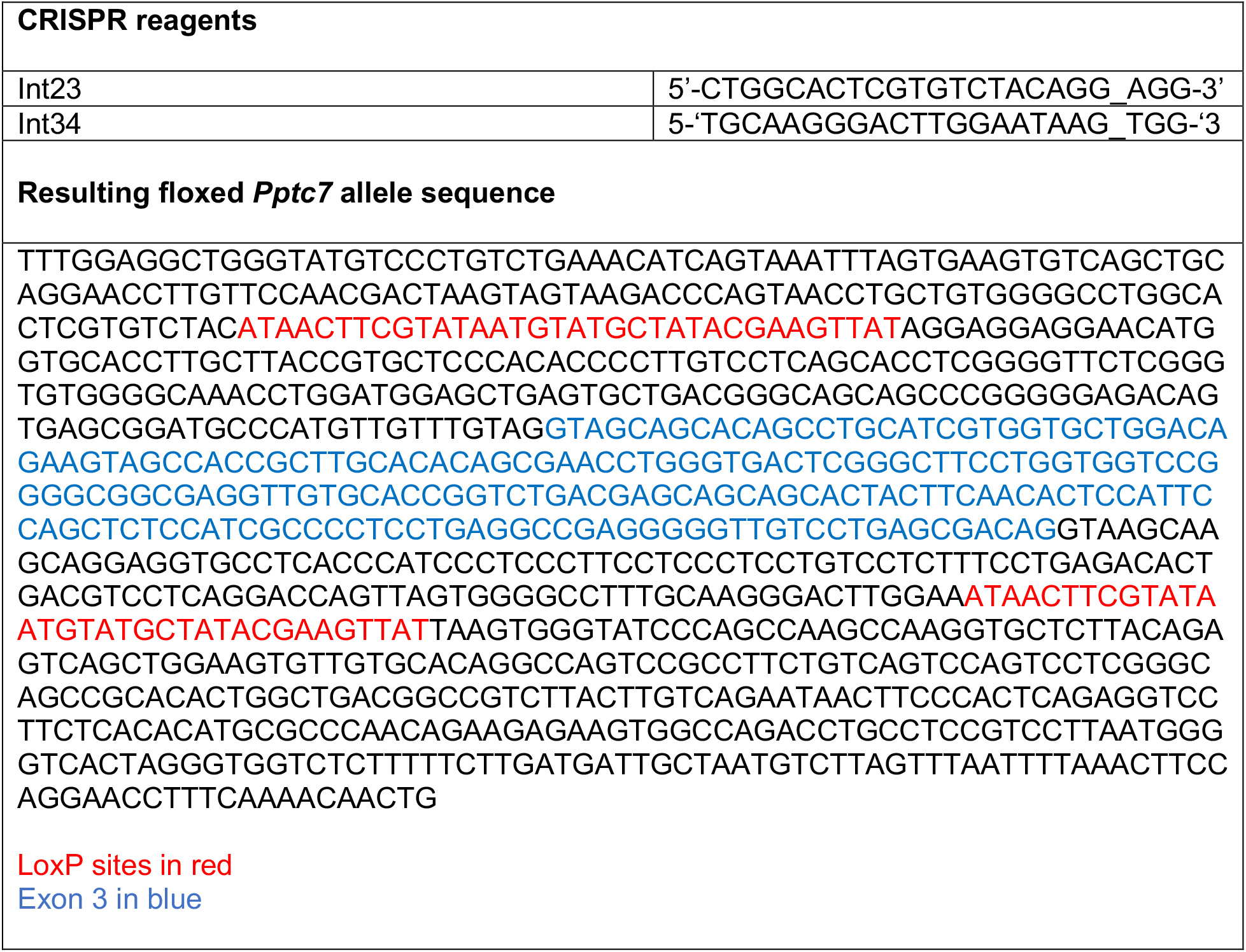
Niemi et al. *Pptc7 maintains mitochondrial protein content by suppressing receptor-mediated mitophagy* Sequences of CRISPR reagents used to generate the *Pptc7* conditional mouse (Int23 and Int24) and the sequence of the resulting floxed allele found in the *Pptc7* conditional mouse model are found below.

**Supplementary Table 2.**
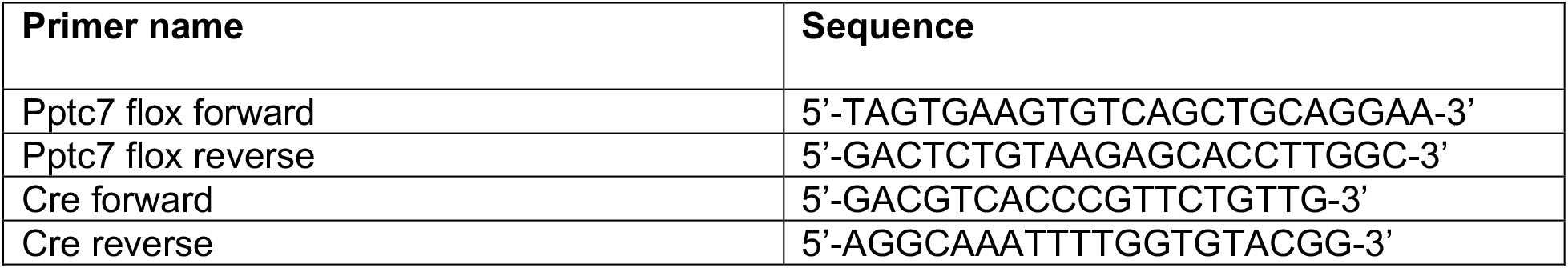
Niemi et al. *Pptc7 maintains mitochondrial protein content by suppressing receptor-mediated mitophagy* Sequences of primers used to genotype the *Pptc7* conditional mouse. Sequences amplify wild type or floxed allele (as full length or excised/knockout) and the UBC-Cre-ER^T2^ recombinase used in this study.

**Supplementary Table 3.**
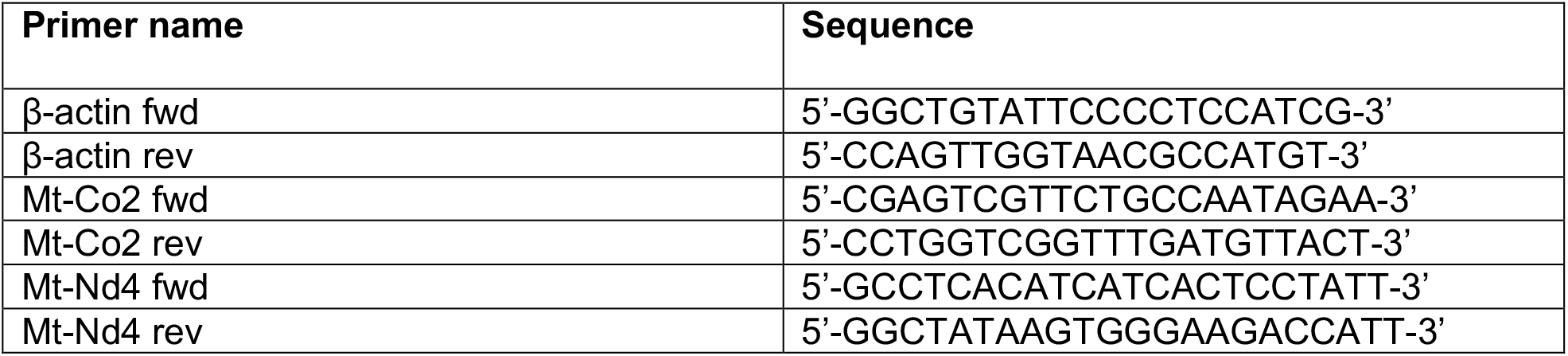
Niemi et al. *Pptc7 maintains mitochondrial protein content by suppressing receptor-mediated mitophagy* Sequences of primers used to amplify nuclear DNA (β-actin) and mitochondrial DNA (mtDNA, Mt-Co2 and Mt-Nd4).

**Supplementary Table 4.**
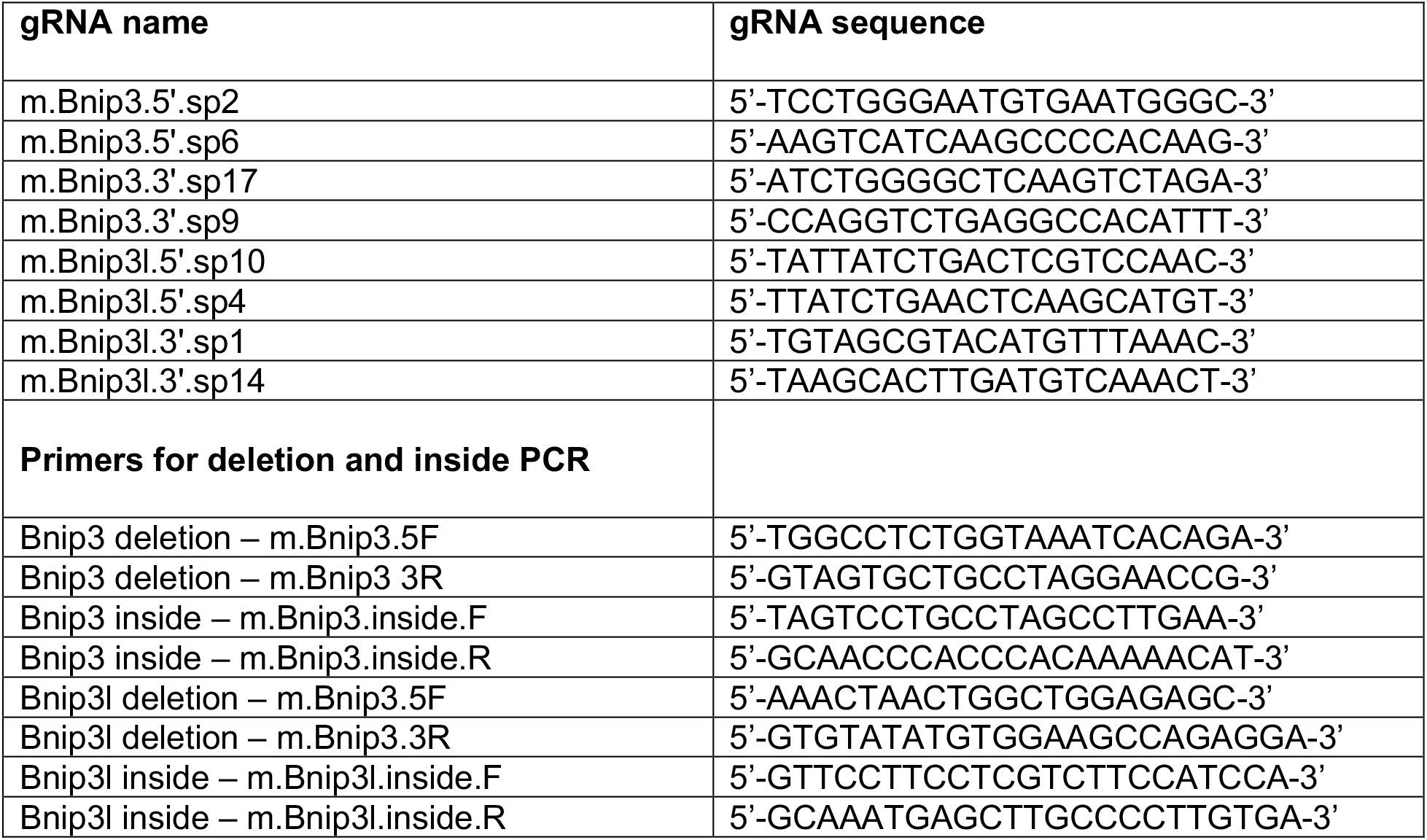
Niemi et al. *Pptc7 maintains mitochondrial protein content by suppressing receptor-mediated mitophagy* Sequences guide RNAs (gRNAs) and primers for detecting *Bnip3* and *Bnip3l* deletion products post-CRISPR.

## References

Alsina, D., Lytovchenko, O., Schab, A., Atanassov, I., Schober, F. A., Jiang, M., Koolmeister, C., Wedell, A., Taylor, R. W., Wredenberg, A., & Larsson, N. (2020). FBXL4 deficiency increases mitochondrial removal by autophagy. EMBO Molecular Medicine. https://doi.org/10.15252/emmm.201911659

Aoki, Y., Kanki, T., Hirota, Y., Kurihara, Y., Saigusa, T., Uchiumi, T., & Kang, D. (2011). Phosphorylation of Serine 114 on Atg32 mediates mitophagy. Molecular Biology of the Cell, 22(17), 3206–3217. https://doi.org/10.1091/mbc.e11-02-0145

Calvo, S. E., Clauser, K. R., & Mootha, V. K. (2016). MitoCarta2.0: An updated inventory of mammalian mitochondrial proteins. Nucleic Acids Research, 44(D1), D1251–D1257. https://doi.org/10.1093/nar/gkv1003

Chen, G., Han, Z., Feng, D., Chen, Y., Chen, L., Wu, H., Huang, L., Zhou, C., Cai, X., Fu, C., Duan, L., Wang, X., Liu, L., Liu, X., Shen, Y., Zhu, Y., & Chen, Q. (2014). A Regulatory Signaling Loop Comprising the PGAM5 Phosphatase and CK2 Controls Receptor-Mediated Mitophagy. Molecular Cell, 54(3), 362–377. https://doi.org/10.1016/j.molcel.2014.02.034

Chung, J., Wittig, J. G., Ghamari, A., Maeda, M., Dailey, T. A., Bergonia, H., Kafina, M. D., Coughlin, E. E., Minogue, C. E., Hebert, A. S., Li, L., Kaplan, J., Lodish, H. F., Bauer, D. E., Orkin, S. H., Cantor, A. B., Maeda, T., Phillips, J. D., Coon, J. J., … Paw, B. H. (2017, May 29). Erythropoietin signaling regulates heme biosynthesis. ELife; eLife Sciences Publications Limited. https://doi.org/10.7554/eLife.24767

da Silva Rosa, S. C., Martens, M. D., Field, J. T., Nguyen, L., Kereliuk, S. M., Hai, Y., Chapman, D., Diehl-Jones, W., Aliani, M., West, A. R., Thliveris, J., Ghavami, S., Rampitsch, C., Dolinsky, V. W., & Gordon, J. W. (2021). BNIP3L/Nix-induced mitochondrial fission, mitophagy, and impaired myocyte glucose uptake are abrogated by PRKA/PKA phosphorylation. Autophagy, 17(9), 2257–2272. https://doi.org/10.1080/15548627.2020.1821548

Elcocks, H., Brazel, A. J., McCarron, K. R., Kaulich, M., Husnjak, K., Mortiboys, H., Clague, M. J., & Urbé, S. (2022). FBXL4 deficiency promotes mitophagy by elevating NIX [Preprint]. Cell Biology. https://doi.org/10.1101/2022.10.11.511735

Floyd, B. J., Wilkerson, E. M., Veling, M. T., Minogue, C. E., Xia, C., Beebe, E. T., Wrobel, R. L., Cho, H., Kremer, L. S., Alston, C. L., Gromek, K. A., Dolan, B. K., Ulbrich, A., Stefely, J. A., Bohl, S. L., Werner, K. M., Jochem, A., Westphall, M. S., Rensvold, J. W., … Pagliarini, D. J. (2016). Mitochondrial Protein Interaction Mapping Identifies Regulators of Respiratory Chain Function. Molecular Cell, 63(4), 621–632. https://doi.org/10.1016/j.molcel.2016.06.033

Gok, M. O., Connor, O. M., Wang, X., Menezes, C. J., Llamas, C. B., Mishra, P., & Friedman, J. R. (2023). The outer mitochondrial membrane protein TMEM11 demarcates spatially restricted BNIP3/BNIP3L-mediated mitophagy. Journal of Cell Biology, 222(4), e202204021. https://doi.org/10.1083/jcb.202204021

Grimsrud, P. A., Carson, J. J., Hebert, A. S., Hubler, S. L., Niemi, N. M., Bailey, D. J., Jochem, A., Stapleton, D. S., Keller, M. P., Westphall, M. S., Yandell, B. S., Attie, A. D., Coon, J. J., & Pagliarini, D. J. (2012). A quantitative map of the liver mitochondrial phosphoproteome reveals posttranslational control of ketogenesis. Cell Metabolism, 16(5), 672–683. https://doi.org/10.1016/j.cmet.2012.10.004

Guo, X., Niemi, N. M., Hutchins, P. D., Condon, S. G. F., Jochem, A., Ulbrich, A., Higbee, A. J., Russell, J. D., Senes, A., Coon, J. J., & Pagliarini, D. J. (2017). Ptc7p Dephosphorylates Select Mitochondrial Proteins to Enhance Metabolic Function. Cell Reports, 18(2), 307–313. https://doi.org/10.1016/j.celrep.2016.12.049

He, Y.-L., Li, J., Gong, S.-H., Cheng, X., Zhao, M., Cao, Y., Zhao, T., Zhao, Y.-Q., Fan, M., Wu, H.-T., Zhu, L.-L., & Wu, L.-Y. (2022). BNIP3 phosphorylation by JNK1/2 promotes mitophagy via enhancing its stability under hypoxia. Cell Death & Disease, 13(11), Article 11. https://doi.org/10.1038/s41419-022-05418-z

Hung, V., Lam, S. S., Udeshi, N. D., Svinkina, T., Guzman, G., Mootha, V. K., Carr, S. A., & Ting, A. Y. (2017, April 25). Proteomic mapping of cytosol-facing outer mitochondrial and ER membranes in living human cells by proximity biotinylation. ELife. https://doi.org/10.7554/eLife.24463

Huttlin, E. L., Bruckner, R. J., Navarrete-Perea, J., Cannon, J. R., Baltier, K., Gebreab, F., Gygi, M. P., Thornock, A., Zarraga, G., Tam, S., Szpyt, J., Gassaway, B. M., Panov, A., Parzen, H., Fu, S., Golbazi, A., Maenpaa, E., Stricker, K., Guha Thakurta, S., … Gygi, S. P. (2021). Dual proteome-scale networks reveal cell-specific remodeling of the human interactome. Cell, 184(11), 3022-3040.e28. https://doi.org/10.1016/j.cell.2021.04.011

Huttlin, E. L., Bruckner, R. J., Paulo, J. A., Cannon, J. R., Ting, L., Baltier, K., Colby, G., Gebreab, F., Gygi, M. P., Parzen, H., Szpyt, J., Tam, S., Zarraga, G., Pontano-Vaites, L., Swarup, S., White, A. E., Schweppe, D. K., Rad, R., Erickson, B. K., … Harper, J. W. (2017). Architecture of the human interactome defines protein communities and disease networks. Nature, 545(7655), Article 7655. https://doi.org/10.1038/nature22366

Huttlin, E. L., Ting, L., Bruckner, R. J., Gebreab, F., Gygi, M. P., Szpyt, J., Tam, S., Zarraga, G., Colby, G., Baltier, K., Dong, R., Guarani, V., Vaites, L. P., Ordureau, A., Rad, R., Erickson, B. K., Wühr, M., Chick, J., Zhai, B., … Gygi, S. P. (2015). The BioPlex Network: A Systematic Exploration of the Human Interactome. Cell, 162(2), 425–440. https://doi.org/10.1016/j.cell.2015.06.043

Jiang, H., & Cao, Y. (2022). A mitochondrial SCF-FBXL4 ubiquitin E3 ligase complex restrains excessive mitophagy to prevent mitochondrial disease [Preprint]. Cell Biology. https://doi.org/10.1101/2022.11.11.516094

Juneau, K., Nislow, C., & Davis, R. W. (2009). Alternative Splicing of PTC7 in Saccharomyces cerevisiae Determines Protein Localization. Genetics, 183(1), 185–194. https://doi.org/10.1534/genetics.109.105155

Kanki, T., Kurihara, Y., Jin, X., Goda, T., Ono, Y., Aihara, M., Hirota, Y., Saigusa, T., Aoki, Y., Uchiumi, T., & Kang, D. (2013). Casein kinase 2 is essential for mitophagy. EMBO Reports, 14(9), 788–794. https://doi.org/10.1038/embor.2013.114

Kravic, B., Harbauer, A. B., Romanello, V., Simeone, L., Vögtle, F.-N., Kaiser, T., Straubinger, M., Huraskin, D., Böttcher, M., Cerqua, C., Martin, E. D., Poveda-Huertes, D., Buttgereit, A., Rabalski, A. J., Heuss, D., Rudolf, R., Friedrich, O., Litchfield, D., Marber, M., … Hashemolhosseini, S. (2018). In mammalian skeletal muscle, phosphorylation of TOMM22 by protein kinase CSNK2/CK2 controls mitophagy. Autophagy, 14(2), 311–335. https://doi.org/10.1080/15548627.2017.1403716

Liu, K. E., & Frazier, W. A. (2015). Phosphorylation of the BNIP3 C-Terminus Inhibits Mitochondrial Damage and Cell Death without Blocking Autophagy. PLOS ONE, 10(6), e0129667. https://doi.org/10.1371/journal.pone.0129667

Mann, M., Hansen, F. M., Kremer, L. S., Karayel, O., Kühl, I., Bludau, I., & Larsson, N.-G. (2022). Mitochondrial phosphoproteomes are functionally specialized across tissues (p. 2022.03.23.485457). bioRxiv. https://doi.org/10.1101/2022.03.23.485457

Martín-Montalvo, A., González-Mariscal, I., Pomares-Viciana, T., Padilla-López, S., Ballesteros, M., Vazquez-Fonseca, L., Gandolfo, P., Brautigan, D. L., Navas, P., & Santos-Ocaña, C. (2013). The Phosphatase Ptc7 Induces Coenzyme Q Biosynthesis by Activating the Hydroxylase Coq7 in Yeast *. Journal of Biological Chemistry, 288(39), 28126–28137. https://doi.org/10.1074/jbc.M113.474494

Murakawa, T., Yamaguchi, O., Hashimoto, A., Hikoso, S., Takeda, T., Oka, T., Yasui, H., Ueda, H., Akazawa, Y., Nakayama, H., Taneike, M., Misaka, T., Omiya, S., Shah, A. M., Yamamoto, A., Nishida, K., Ohsumi, Y., Okamoto, K., Sakata, Y., & Otsu, K. (2015). Bcl-2-like protein 13 is a mammalian Atg32 homologue that mediates mitophagy and mitochondrial fragmentation. Nature Communications, 6(1), Article 1. https://doi.org/10.1038/ncomms8527

Nguyen-Dien, G. T., Kozul, K.-L., Cui, Y., Townsend, B., Kulkarni, P. G., Ooi, S. S., Marzio, A., Carrodus, N., Zuryn, S., Pagano, M., Parton, R. G., Lazarou, M., Millard, S., Taylor, R. W., Collins, B. M., Jones, M. J. K., & Pagan, J. K. (2022). FBXL4 suppresses mitophagy by restricting the accumulation of NIX and BNIP3 mitophagy receptors (p. 2022.10.12.511867). bioRxiv. https://doi.org/10.1101/2022.10.12.511867

Niemi, N. M., & MacKeigan, J. P. (2013). Mitochondrial Phosphorylation in Apoptosis: Flipping the Death Switch. Antioxidants & Redox Signaling, 19(6), 572–582. https://doi.org/10.1089/ars.2012.4982

Niemi, N. M., & Pagliarini, D. J. (2021). The extensive and functionally uncharacterized mitochondrial phosphoproteome. Journal of Biological Chemistry, 297(1). https://doi.org/10.1016/j.jbc.2021.100880

Niemi, N. M., Wilson, G. M., Overmyer, K. A., Vögtle, F.-N., Myketin, L., Lohman, D. C., Schueler, K. L., Attie, A. D., Meisinger, C., Coon, J. J., & Pagliarini, D. J. (2019). Pptc7 is an essential phosphatase for promoting mammalian mitochondrial metabolism and biogenesis. Nature Communications, 10(1), 3197. https://doi.org/10.1038/s41467-019-11047-6

Pagliarini, D. J., Calvo, S. E., Chang, B., Sheth, S. A., Vafai, S. B., Ong, S.-E., Walford, G. A., Sugiana, C., Boneh, A., Chen, W. K., Hill, D. E., Vidal, M., Evans, J. G., Thorburn, D. R., Carr, S. A., & Mootha, V. K. (2008). A Mitochondrial Protein Compendium Elucidates Complex I Disease Biology. Cell, 134(1), 112–123. https://doi.org/10.1016/j.cell.2008.06.016

Pickles, S., Vigié, P., & Youle, R. J. (2018). Mitophagy and Quality Control Mechanisms in Mitochondrial Maintenance. Current Biology, 28(4), R170–R185. https://doi.org/10.1016/j.cub.2018.01.004

Poole, L. P., Bock-Hughes, A., Berardi, D. E., & Macleod, K. F. (2021). ULK1 promotes mitophagy via phosphorylation and stabilization of BNIP3. Scientific Reports, 11(1), Article 1. https://doi.org/10.1038/s41598-021-00170-4

Quadros, R. M., Miura, H., Harms, D. W., Akatsuka, H., Sato, T., Aida, T., Redder, R., Richardson, G. P., Inagaki, Y., Sakai, D., Buckley, S. M., Seshacharyulu, P., Batra, S. K., Behlke, M. A., Zeiner, S. A., Jacobi, A. M., Izu, Y., Thoreson, W. B., Urness, L. D., … Gurumurthy, C. B. (2017). Easi-CRISPR: A robust method for one-step generation of mice carrying conditional and insertion alleles using long ssDNA donors and CRISPR ribonucleoproteins. Genome Biology, 18(1), 92. https://doi.org/10.1186/s13059-017-1220-4

Rath, S., Sharma, R., Gupta, R., Ast, T., Chan, C., Durham, T. J., Goodman, R. P., Grabarek, Z., Haas, M. E., Hung, W. H. W., Joshi, P. R., Jourdain, A. A., Kim, S. H., Kotrys, A. V., Lam, S. S., McCoy, J. G., Meisel, J. D., Miranda, M., Panda, A., … Mootha, V. K. (2021). MitoCarta3.0: An updated mitochondrial proteome now with sub-organelle localization and pathway annotations. Nucleic Acids Research, 49(D1), D1541–D1547. https://doi.org/10.1093/nar/gkaa1011

Rensvold, J. W., Shishkova, E., Sverchkov, Y., Miller, I. J., Cetinkaya, A., Pyle, A., Manicki, M., Brademan, D. R., Alanay, Y., Raiman, J., Jochem, A., Hutchins, P. D., Peters, S. R., Linke, V., Overmyer, K. A., Salome, A. Z., Hebert, A. S., Vincent, C. E., Kwiecien, N. W., … Pagliarini, D. J. (2022). Defining mitochondrial protein functions through deep multiomic profiling. Nature, 1–7. https://doi.org/10.1038/s41586-022-04765-3

Rhee, H.-W., Zou, P., Udeshi, N. D., Martell, J. D., Mootha, V. K., Carr, S. A., & Ting, A. Y. (2013). Proteomic Mapping of Mitochondria in Living Cells via Spatially Restricted Enzymatic Tagging. Science, 339(6125), 1328–1331. https://doi.org/10.1126/science.1230593

Rogov, V. V., Suzuki, H., Marinković, M., Lang, V., Kato, R., Kawasaki, M., Buljubašić, M., Šprung, M., Rogova, N., Wakatsuki, S., Hamacher-Brady, A., Dötsch, V., Dikic, I., Brady, N. R., & Novak, I. (2017). Phosphorylation of the mitochondrial autophagy receptor Nix enhances its interaction with LC3 proteins. Scientific Reports, 7(1), 1131. https://doi.org/10.1038/s41598-017-01258-6

Ruzankina, Y., Pinzon-Guzman, C., Asare, A., Ong, T., Pontano, L., Cotsarelis, G., Zediak, V. P., Velez, M., Bhandoola, A., & Brown, E. J. (2007). Deletion of the Developmentally Essential Gene ATR in Adult Mice Leads to Age-Related Phenotypes and Stem Cell Loss. Cell Stem Cell, 1(1), 113–126. https://doi.org/10.1016/j.stem.2007.03.002

Schäfer, J. A., Bozkurt, S., Michaelis, J. B., Klann, K., & Münch, C. (2022). Global mitochondrial protein import proteomics reveal distinct regulation by translation and translocation machinery. Molecular Cell, 82(2), 435-446.e7. https://doi.org/10.1016/j.molcel.2021.11.004

Song, J., Herrmann, J. M., & Becker, T. (2021). Quality control of the mitochondrial proteome. Nature Reviews Molecular Cell Biology, 22(1), Article 1. https://doi.org/10.1038/s41580-020-00300-2

Sun, N., Malide, D., Liu, J., Rovira, I. I., Combs, C. A., & Finkel, T. (2017). A fluorescence-based imaging method to measure in vitro and in vivo mitophagy using mt-Keima. Nature Protocols, 12(8), Article 8. https://doi.org/10.1038/nprot.2017.060

Walter, C., Marada, A., Suhm, T., Ernsberger, R., Muders, V., Kücükköse, C., Sánchez-Martín, P., Hu, Z., Aich, A., Loroch, S., Solari, F. A., Poveda-Huertes, D., Schwierzok, A., Pommerening, H., Matic, S., Brix, J., Sickmann, A., Kraft, C., Dengjel, J., … Meisinger, C. (2021). Global kinome profiling reveals DYRK1A as critical activator of the human mitochondrial import machinery. Nature Communications, 12(1), 4284. https://doi.org/10.1038/s41467-021-24426-9

Wang, L., Qi, H., Tang, Y., & Shen, H.-M. (2020). Post-translational Modifications of Key Machinery in the Control of Mitophagy. Trends in Biochemical Sciences, 45(1), 58–75. https://doi.org/10.1016/j.tibs.2019.08.002

Williams, C. C., Jan, C. H., & Weissman, J. S. (2014). Targeting and plasticity of mitochondrial proteins revealed by proximity-specific ribosome profiling. Science, 346(6210), 748–751. https://doi.org/10.1126/science.1257522

Zhao, J.-F., Rodger, C. E., Allen, G. F. G., Weidlich, S., & Ganley, I. G. (2020). HIF1α-dependent mitophagy facilitates cardiomyoblast differentiation. Cell Stress, 4(5), 99–113. https://doi.org/10.15698/cst2020.05.220

Zhu, Y., Massen, S., Terenzio, M., Lang, V., Chen-Lindner, S., Eils, R., Novak, I., Dikic, I., Hamacher-Brady, A., & Brady, N. R. (2013). Modulation of Serines 17 and 24 in the LC3-interacting Region of Bnip3 Determines Pro-survival Mitophagy versus Apoptosis. Journal of Biological Chemistry, 288(2), 1099–1113. https://doi.org/10.1074/jbc.M112.399345

